# Bridging Light and Sound: a Spironaphtopyran-Rhodamine Dyad with High-Contrast Photoswitching Between Fluorescence and Photoacoustic Signal

**DOI:** 10.1101/2025.10.17.683093

**Authors:** Nikita Kaydanov, Magdalena Olesińska-Mönch, Morgane Leite, Robert Prevedel, Claire Deo

## Abstract

Fluorescence and photoacoustic imaging are complementary modalities that provide distinct advantages for biological imaging: fluorescence microscopy offers high sensitivity and resolution, while photoacoustic imaging enables deeper penetration in complex tissue. Leveraging the strengths of both modalities through optically switchable contrast agents can offer enhanced imaging contrast and facilitate dual-modality imaging. Here, we report a photoswitchable probe capable of toggling between high fluorescence and high photoacoustic signal upon illumination, exploiting Förster Resonance Energy Transfer (FRET). We engineer novel spironaphtopyran photoswitches which undergo reversible photoisomerization between absorbing and non-absorbing states. Their photoswitching properties were systematically characterized, establishing structure-properties relationships, and providing the first photoacoustic investigation into this class of compounds. The best-performing switch was incorporated into a FRET dyad with a rhodamine fluorophore, which exhibits robust, reversible switching between fluorescent and photoacoustic-dominant states with excellent contrast *in vitro*, establishing a foundation for multimodal imaging probes with promising potential for dynamic correlative imaging.

## 1. INTRODUCTION

Optical imaging modalities are powerful tools for the study of biological systems. Fluorescence (Fl) microscopy offers excellent sensitivity and spatial resolution but is fundamentally limited in imaging depth due to light scattering.[1] In contrast, photoacoustic (PA) imaging, which combines optical excitation with ultrasound detection, enables imaging several centimeters deep in complex biological tissues.[2, 3] Critically, the performance of these techniques depends on the availability of robust contrast agents, able to generate specific signal. Among them, photoswitchable reporters which contrast can be modulated using light offer unique advantages. Indeed, such reporters have enabled breakthroughs in super-resolution fluorescence microscopy,[4, 5] facilitated background corrections in photoacoustic imaging,[6-8] and have been exploited in the design of photoresponsive functional tools such as biosensors.[9-11]

Extending this concept to dual-modality probes that can be optically toggled between fluorescence- and photoacoustic-dominant states can offer complementary information across scales and depths, which presents exciting opportunities for correlative imaging ranging from neuroimaging in model organisms to potential clinical applications.[12-15] However, achieving high contrast in both modalities is challenging, as it requires precise control over how absorbed energy is distributed between radiative (fluorescence) and non-radiative (photoacoustic) decay pathways, which are inherently competitive. We reasoned that this could be achieved by exploiting Förster Resonance Energy Transfer (FRET) between a photochromic acceptor and a fluorescent donor (Figure 1). In the dark state, efficient FRET between the two moieties would lead primarily to non-radiative relaxation, suppressing fluorescence, while retaining strong PA signal due to the high absorbance of both donor and acceptor. Upon illumination, photoisomerization to the non-absorbing isomer would abolish FRET, resulting in a large fluorescence increase and concomitant photoacoustic signal decrease, therefore efficiently modulating contrast across both modalities. To engineer such a FRET-based molecular reporter, a key requirement is an efficient photoswitchable acceptor which absorbs strongly in the dark state but becomes transparent following light activation (i.e. a “negative” photoswitch[16]). For this purpose, we focused on spiropyran derivatives, which represent a versatile class of photochromic compounds capable of reversible interconversion between a non-absorbing spiro (SP) form and a strongly absorbing merocyanine (MC) form.[17-20] Importantly, depending on their structural features, derivatives of this family have been found to be stabilized as the MC isomer in the dark state,[21, 22] which predispose them to negative photoswitching behavior, and have additionally been exploited as FRET acceptors with common fluorophores,[10, 23, 24] which makes them ideal candidates for our approach.

**Figure 1.**
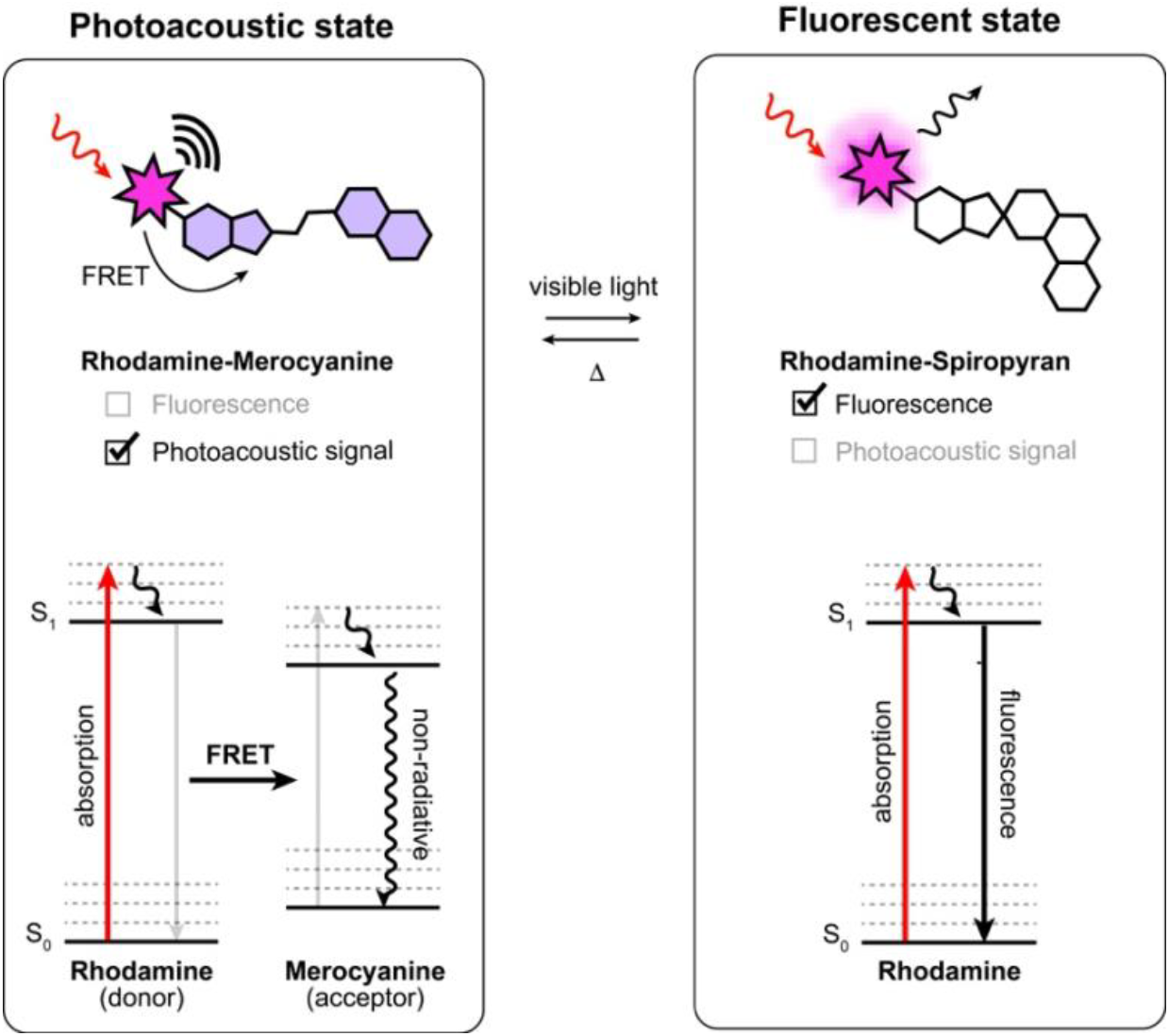
General approach for the design of dual-modality fluorescent/photoacoustic photoswitchable probes based on FRET between a rhodamine fluorophore and a spiropyran (SP)-merocyanine (MC) photoswitch. S_0_ and S1 denote the ground state and first electronic excited state of the fluorophore, respectively.

Here, we design and synthesize a library of spironaphthopyran derivatives and systematically characterize their photoswitching properties in absorption and photoacoustic signal using a custom-built multimodal spectroscopy platform. To the best of our knowledge, this constitutes the first photoacoustic investigation into this class of compounds. We identify a robust negative photoswitch that undergoes rapid and efficient isomerization under visible light. Coupling this photoswitch with a rhodamine fluorophore yields a photoswitchable dyad that toggles reversibly between PA-dominant and FL-dominant states with fast kinetics and large dynamic range. This system provides dynamic dual-modality contrast within a single molecular probe, opening future opportunities for deep-tissue and high-resolution imaging.

## 2. RESULTS AND DISCUSSION

### 2.1 Design and synthesis

To develop a negative photoswitch suitable for FRET, we focused on the spironaphthopyran scaffold (Figure 2a). These photochromic compounds of the spiropyran family have only been scarcely investigated,[24, 25] but can advantageously offer a red-shift in absorption in the merocyanine form due to their extended conjugation. This can provide improved spectral overlap with fluorophores in the visible range, which is an essential criterion for efficient FRET. In addition, spironaphthopyrans have been shown to be stabilized in their MC form in polar solvents, making them promising candidates for negative photoswitching. We therefore synthesized a new family of spironaphtopyrans bearing various electron-withdrawing substituents at the 6-position, which lies para to the phenolate oxygen of the MC form and is known to play a critical role in the photochromic behavior of spiropyran switches. The substituents included nitro (– NO_2_),[24] halogens (–F, –Br), and carbonyl groups such as methyl ester (–CO_2_Me) and aldehyde (–CHO) (Figure 2a). To enable covalent conjugation to a fluorophore, a carboxylic acid functional group was introduced at the 5′-position of the indolenine ring. The target compounds were synthesized using established methods (Figure S1). Specifically, Duff formylation of commercially available naphthol derivatives afforded substituted naphthaldehydes, which then underwent Knoevenagel condensation with the Fischer base of indolenine **S6** to provide spironaphthopyrans **1**–**5**.

**Figure 2.**
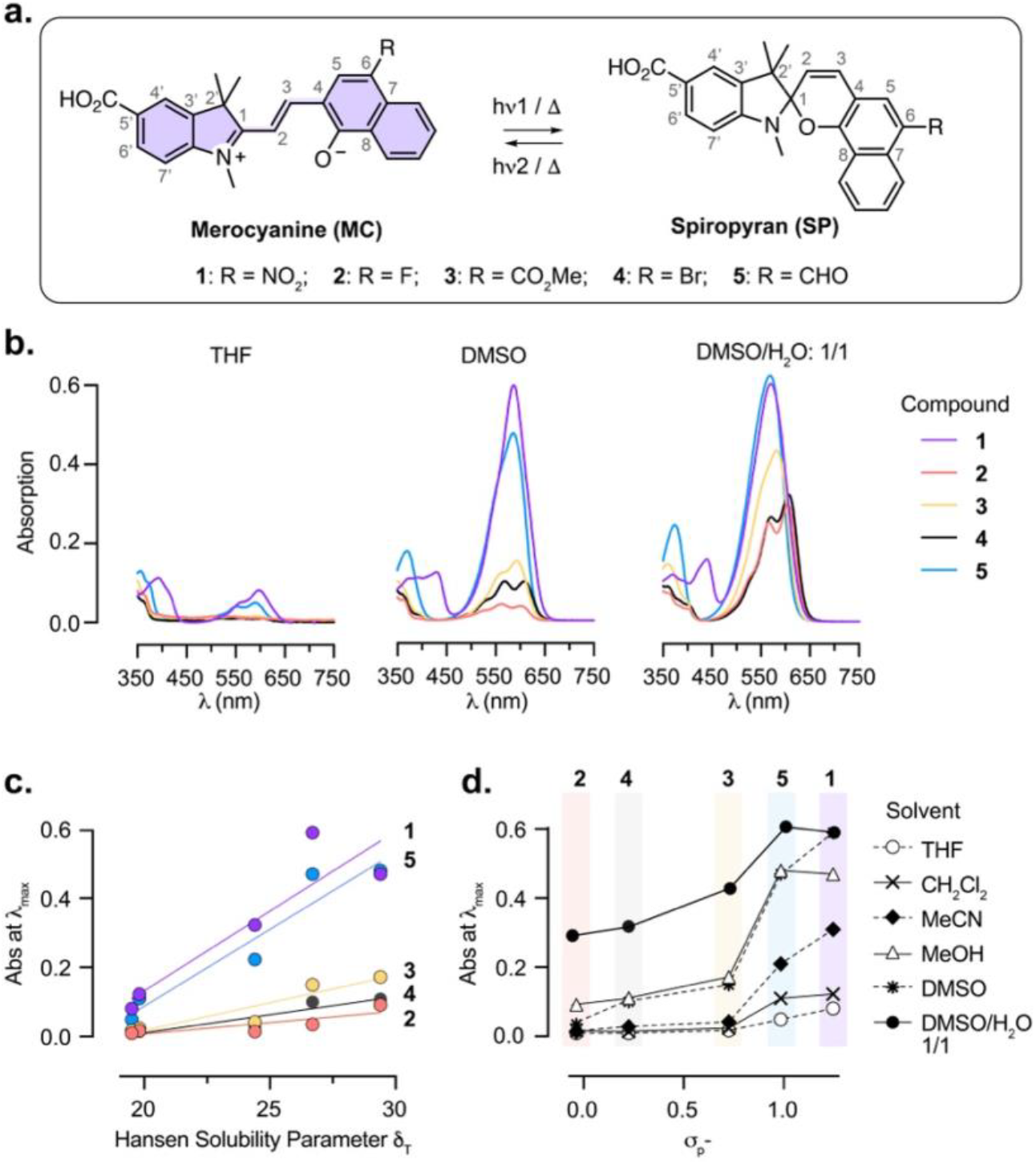
**a**. Structure and photoisomerization of spironaphthopyrans **1**–**5** studied in this work. **b**. Absorption spectra of the dark state of compounds **1**–**5** (10 μM) in THF, DMSO and 1/1 DMSO/H_2_O mixture. **c**. Relationship between the Hansen Solubility Parameter δ_T_ of the solvents used and the absorbance at λ_max_ for compounds **1**–**5** in the dark state. **d**. Relationship between the Hammett constant σ_P_^-^ of the 6-position substituent and the absorbance at λ_max_ for compounds **1**–**5** in different solvents, in the dark state.

### 2.2 Photophysical characterization in the dark state

Spiropyran derivatives exist in equilibrium between the closed, colourless spiro (SP) and the open, highly absorbing merocyanine (MC) isomers in solution, with this equilibrium being highly dependent on the local environment.[17] To evaluate the potential of compounds **1–5** as negative photoswitches (i.e. thermally stable as the MC isomer), we measured their absorption spectra in the dark state in various solvents (Figure 2b, Figure S2). In all cases, the compounds exhibited absorption in the visible range (λ_max_ = 585−609 nm), corresponding to the MC isomers and consistent with their extended conjugation. As is commonly the case for photoswitchable compounds, the MC form of the switches displayed only very weak fluorescence in the visible (λ_em_ = 617−631 nm) with Φ_FL_ ≤ 0.04 in DMSO (Figure S3). In certain solvent–compound combinations, an additional blue-shifted shoulder was observed, which can be attributed to MC isomeric forms, such as the quinoid resonance structure or other conformers.[25, 26] The relative populations of the SP and MC forms (inferred from the intensity of the MC absorption band), varied markedly with solvent polarity. In low-polarity solvents such as tetrahydrofuran (THF), the SP form was favored, whereas in highly polar solvents such as methanol and dimethyl sulfoxide (DMSO), the MC form was increasingly stabilized. This trend correlated with the Hansen solubility parameter (δ_T_) of the solvents,[27] supporting the role of solvent interactions in modulating isomer distribution (Figure 2c). The addition of water further promoted formation of the MC form, as indicated by increased absorption. Notably, the extent of this solvent-induced shift varied among the compounds. Compounds **1** and **5** showed a substantially higher absorption in the visible across all solvents tested, which was relatively unaffected by addition of water, while compounds **2**–**4** exhibited a more pronounced increase in MC absorption upon polarity increase and water addition. This suggests that compounds **1** and **5** display a substantially larger proportion of the MC isomer in the dark state, which was supported by DFT calculations (see Supplementary Note), and quantified by ^1^H NMR in (CD_3_)_2_SO (Figure S4). Compounds **1** and **5** exist predominantly as the MC form in the dark, with compound **1** being the more strongly shifted with 66% of the MC isomer, while compounds **2**–**4** were present at >85% of the SP form. We note that the peaks corresponding to the MC isomer of compounds **3** and **5** are broadened, introducing uncertainty in the integration, which may arise from the presence of multiple MC conformers.[28] Overall, the SP/MC equilibrium correlated with the Hammett σ_P_^-^ constants of the substituent at the 6-position,[29] providing a rational framework to tune the equilibrium position within this family of compounds (Figure 2d).[30] Merocyanines are known to undergo rapid hydrolysis in the presence of water, which impairs their broad applicability in biological systems.[31, 32] To investigate the potential of these new compounds for future biological applications, we measured the hydrolysis rates of compounds **1**–**5** in water (Figure S5). Gratifyingly, these novel spironaphtopyrans showed remarkable stability in water compared to the parent nitro-spiropyran **6**, with hydrolysis rate for compounds **1**–**5** also closely correlated with the Hammett σ_P_^-^ constants of the electron-withdrawing substituents. Overall, this suggests that these spironapthopyrans present sufficient stability in aqueous solution for biological applications.

### 2.3 Photoisomerization

Next, we investigated the photoisomerization behavior of compounds **1**–**5**, focusing on the light-induced conversion of the merocyanine (MC) to spiro (SP) form under illumination with visible light. A previous study reported that compound **1** did not exhibit appreciable photoswitching,[24] which we hypothesized to be the result of rapid thermal relaxation from the SP back to the thermodynamically favored MC state. To enable precise characterization with high temporal resolution, we developed a custom-built multimodal optical platform capable of simultaneous absorption (Abs), photoacoustic (PA), and fluorescence (FL) measurements (Figure S6). Illumination was performed using the photoacoustic excitation laser tuned to the absorption maximum (λ_max_) for each compound, while absorbance and photoacoustic signal were monitored during alternating illumination/dark cycles (Figure 3). The duration of the ON and OFF cycles was adjusted for each compound, to capture complete switching kinetics while minimizing photobleaching. All measurements were conducted in anhydrous dimethyl sulfoxide (DMSO), selected for its high polarity (favoring MC stabilization), aprotic nature (preventing protonation effects), and high boiling point (reducing evaporation-related concentration change). Compound concentrations were adjusted to maintain an optical density of 0.1 at λ_max_, ensuring comparable concentration of the MC isomer across all compounds. All five spironaphtopyrans displayed photoswitching which was fully reversible in the dark, albeit with drastically distinct behaviors (Figure 3, Table 1). Importantly, Abs and PA signal exhibited excellent agreement in both amplitude and switching kinetics for all compounds, supporting a strictly absorption-driven switching mechanism (Figure 3). Switching amplitude varied widely among compounds. Compounds **1** and **2** showed near-complete isomerization, with minimal residual visible absorbance at the photostationary state, while compounds **3**–**5** exhibited smaller switching amplitudes (18–28%). Compounds **1, 2, 4**, and **5** exhibited the expected negative switching (i.e., a decrease in Abs and PA signal upon illumination), consistent with MC-to-SP conversion. Interestingly, compound **3** exhibited an unexpected positive switching response, characterized by increased absorbance during illumination. A similar observation was made for compound **5** under prolonged light exposure, where the directionality of the switching inverted as a result of photobleaching, transitioning from turn-off to turn-on (Figure S7). This behavior evidences a more complex photoswitching process involving the formation of an additional metastable species absorbing in the visible range. In anhydrous DMSO here, formation of the protonated form of the merocyanine would be negligible.[26, 33] Therefore, we hypothesize that this third species likely arises from trans-to-cis isomerization, as cis-MC is a well-known intermediate formed during the photoisomerization process (Figure S8).[34-37] The occurrence of this effect only in the carbonyl-substituted compounds (**3** and **5**) may reflect specificity in the spectral features of the cis-MC isomer, as well as important differences in kinetics rates, where the photoconversion to the SP isomer would be unfavorable. Unfortunately, the fast relaxation kinetics prevent further experimental investigation into the nature of the species involved at the photostationnary state, and attempts to identify photobleaching products of compound **5** by HPLC-MS were unsuccessful, showing a complex mixture of numerous products.

**Table 1:** Spectroscopic properties of compounds studied in this work in anhydrous DMSO. λ_max_, λ_em_ and Φ_F_ correspond to the MC isomer. ^a^Extinction coefficient of the dark state mixture, ^b^SP/MC ratio in the dark state. Values are reported as mean ±SEM.

| Compound | $\lambda_{\text{max}}$ (nm) | $\lambda_{\text{em}}$ (nm) | $\Phi_{\text{FL}}$ | $\epsilon_{\text{OFF}}$ ($\text{M}^{-1}\cdot\text{cm}^{-1}$)<sup>a</sup> | SP/MC<sub>OFF</sub><sup>b</sup> | Abs<sub>ON</sub>/Abs<sub>OFF</sub> | PA<sub>ON</sub>/PA<sub>OFF</sub> | FL<sub>ON</sub>/FL<sub>OFF</sub> | $\tau_{\text{ON}}$ (s) | $\tau_{\text{OFF}}$ (s) |
| --- | --- | --- | --- | --- | --- | --- | --- | --- | --- | --- |
| **1** | 586 | 622 | 0.04 | 59000 | 34/66 | 0.05 ± 0.01 | 0.05 ± 0.01 | n.m | 1.26 ± 0.14 (1.26 ± 0.09) | 8.21 ± 0.17 |
| **2** | 560, 603 | 625 | n.m | 4000, 3500 | 86/14 | 0.13 ± 0.01 | 0.09 ± 0.01 | n.m | 0.31 ± 0.01 (0.34 ± 0.02) | 682 ± 200 |
| **3** | 593 | 621 | 0.01 | 15000 | 89/11 | 1.19 ± 0.01 | 1.15 ± 0.01 | n.m | 3.30 ± 0.12 (3.24 ± 0.19) | 2.80 ± 0.10 |
| **4** | 567, 609 | 631 | 0.04 | 9800, 10000 | 89/11 | 0.72 ± 0.01 | 0.63 ± 0.01 | n.m | 0.71 ± 0.01 (0.76 ± 0.01) | 43 ± 9 |
| **5** | 585 | 617 | 0.02 | 47000 | 42/58 | 0.82 ± 0.01 | 0.80 ± 0.02 | n.m | 5.99 ± 0.19 (6.15 ± 0.30) | 5.54 ± 0.13 |
| **Rh** | 569 | 590 | 0.55 | 94000 | - | - | - | - | - | - |
| **Rh-1** | 571 | 594 | 0.05 | 147000 | 23/77 | 0.53 ± 0.01 | 0.41 ± 0.01 | 30.1 ± 1.4 | 1.19 ± 0.04 (1.14 ± 0.03) | 3.72 ± 0.07 |

**Figure 3.**
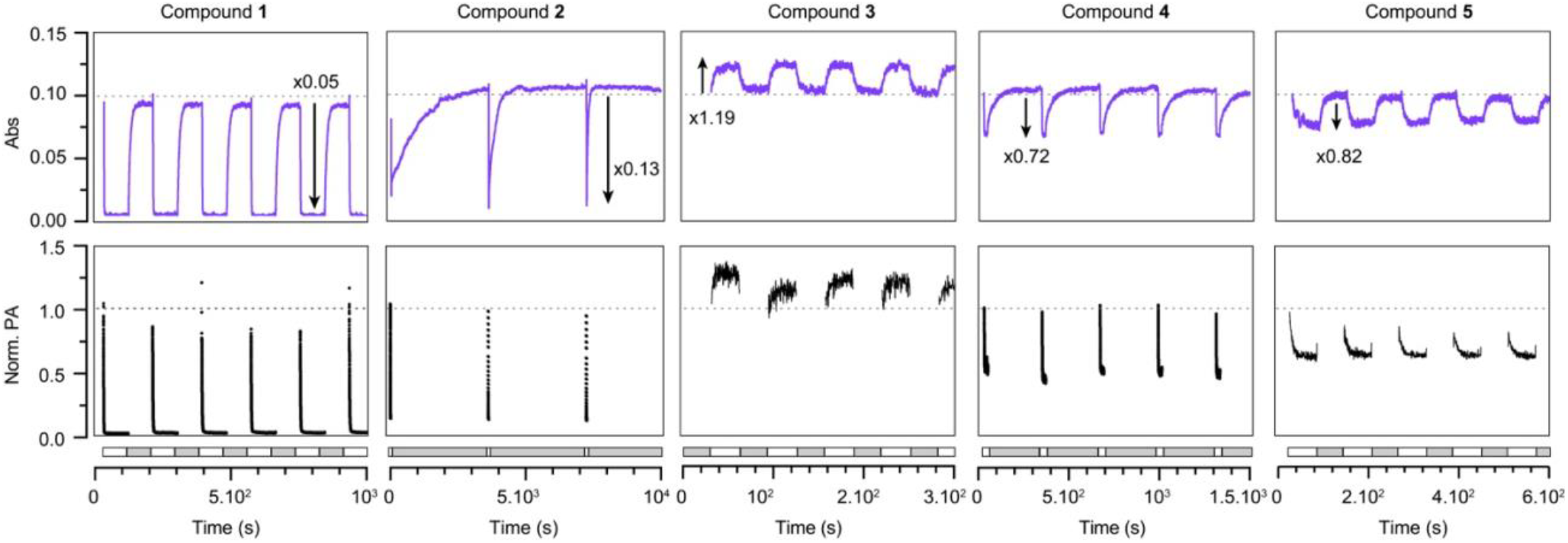
Absorbance and normalized photoacoustic signal of compounds **1-5** upon cycles of illumination and dark. Excitation was performed at λ_max_ for each compound, and absorbance was measured at λ_max_+10 nm. Illumination and dark durations were adjusted for each compound to reach equilibrium and is given as on time/off time: compound **1**: 90s/90s; **2**: 10s/90min; **3**: 30s/30s; **4**: 20s/5min; **5**: 60s/60s. Grey and white boxes underneath correspond to dark and illumination frames, respectively.

Photoswitching and thermal relaxation processes followed first-order kinetics. Under our experimental conditions, photoswitching was rapid (τ_ON_ < 6 s) for all compounds (Table 1, Figure S9). Thermal relaxation kinetics were widely compound-dependent: **1, 3**, and **5** reverted quickly to the MC form, while halogen-substituted **2** and **4** relaxed more slowly. Interestingly, compound **2** displayed progressively faster thermal relaxation over successive switching cycles (Figure 3). We attributed this behavior to increasing water content in the DMSO solution over time, absorbed from the surrounding water bath due to the open cuvette design. Supporting this hypothesis, measurements of the photoswitching behavior of compound **2** in DMSO/water mixtures confirmed that water significantly accelerates SP-to-MC thermal reversion, while the ON-switching process was independent of the presence of water (Figure S10). The addition of water also led to a reduced dynamic range, which could be explained by an increased stabilization of the MC isomer.

### 2.4 Synthesis and characterization of a photoswitchable dyad

Compound **1**, exhibiting the highest proportion of the MC form in the dark state, near-quantitative photoisomerization to the SP form upon illumination, fast switching kinetics, and good photostability, appears as an ideal candidate for integration in a FRET-based multimodal photoswitchable probe. To enable energy transfer-based modulation, compound **1** was conjugated to rhodamine B, chosen as the fluorescent donor due to its high fluorescence quantum yield and excellent fluorescence spectrum overlap with the MC absorption band of **1** (Figure S11). The dyad **Rh-1** was synthesized via amide coupling, with compound **1** appended to the *ortho*-position of rhodamine B with an amide linkage designed to abolish the open-closed equilibrium of the rhodamine,[38] which could interfere with the measurements (Figure S12, Figure 4). As expected, the resulting probe displayed additive absorption features corresponding to both **Rh** and compound **1** (Figure 4b). ^1^H NMR analysis revealed that ∼77% of the dyad existed in the MC form in the dark state in DMSO, higher than for compound **1** alone (Figure S13). This suggests that substitution at the 5′ position influences the SP/MC equilibrium, and can further enhance MC stabilization. In the dark state, dyad **Rh-1** exhibited strongly quenched fluorescence in the visible region (λ_em_ = 594 nm), with Φ_FL_ = 0.05, compared to Φ_FL_ = 0.55 for the parent fluorophore **Rh** (Table 1). This significant quenching indicates highly efficient FRET from **Rh** to the MC form of **1**, with a calculated FRET efficiency >99.9% (see Methods). The residual weak emission observed for **Rh-1** can be attributed to the small proportion of the SP isomer of **Rh-1** at equilibrium in solution, and additionally to the intrinsic low fluorescence of **1** which also contributes to the overall emission spectrum (Figure 4c). Importantly, **Rh-1** exhibited highly efficient photoswitching. Upon illumination, absorption decreased by about 50%, consistent with complete conversion to the SP form, as the residual absorption at the photostationary state can be fully attributed to the **Rh** moiety. As expected, the compound exhibited inverse PA and FL switching. PA signal decreased by ∼60%, slightly larger than the absorbance modulation, which is consistent with the additional contribution of the fluorescence activation further reducing residual PA signal. Fluorescence (monitored at 591 nm which ensures negligible contribution from the fluorescence of **1** itself) increased by an impressive 30-fold upon illumination. As expected, kinetic parameters for the turn-on and turn-off processes were in the same range as parent compound **1** (Table 1, Figure S14). In contrast, rhodamine **Rh** alone did not show any photoswitching behavior, and only photobleaching was observed upon illumination with a loss of 80% of fluorescence over 20 ON/OFF cycles (Figure S15), confirming that the observed signal modulation in **Rh-1** unambiguously stems from MC to SP conversion and concomitant photoswitchable FRET. Nevertheless, photobleaching was also observed for **Rh-1**, suggesting that improvements would be beneficial for future applications (Figure S16). To investigate the possibility to translate **Rh-1** to biological conditions, we examined the effect of water on its photoswitching properties. Water addition substantially increased the photoactivation and thermal relaxation kinetics, but interestingly did not affect the switching amplitude, retaining high contrast (Figure S17). While no photoswitching of **Rh-1** could be measured in fully aqueous conditions, these findings support that photoswitching still occurs efficiently but with kinetics too fast to be measured using our set-up. Altogether, **Rh-1** exhibits dual-modality switching with high dynamic range and fast kinetics, making it a promising scaffold for future imaging applications.

**Figure 4.**
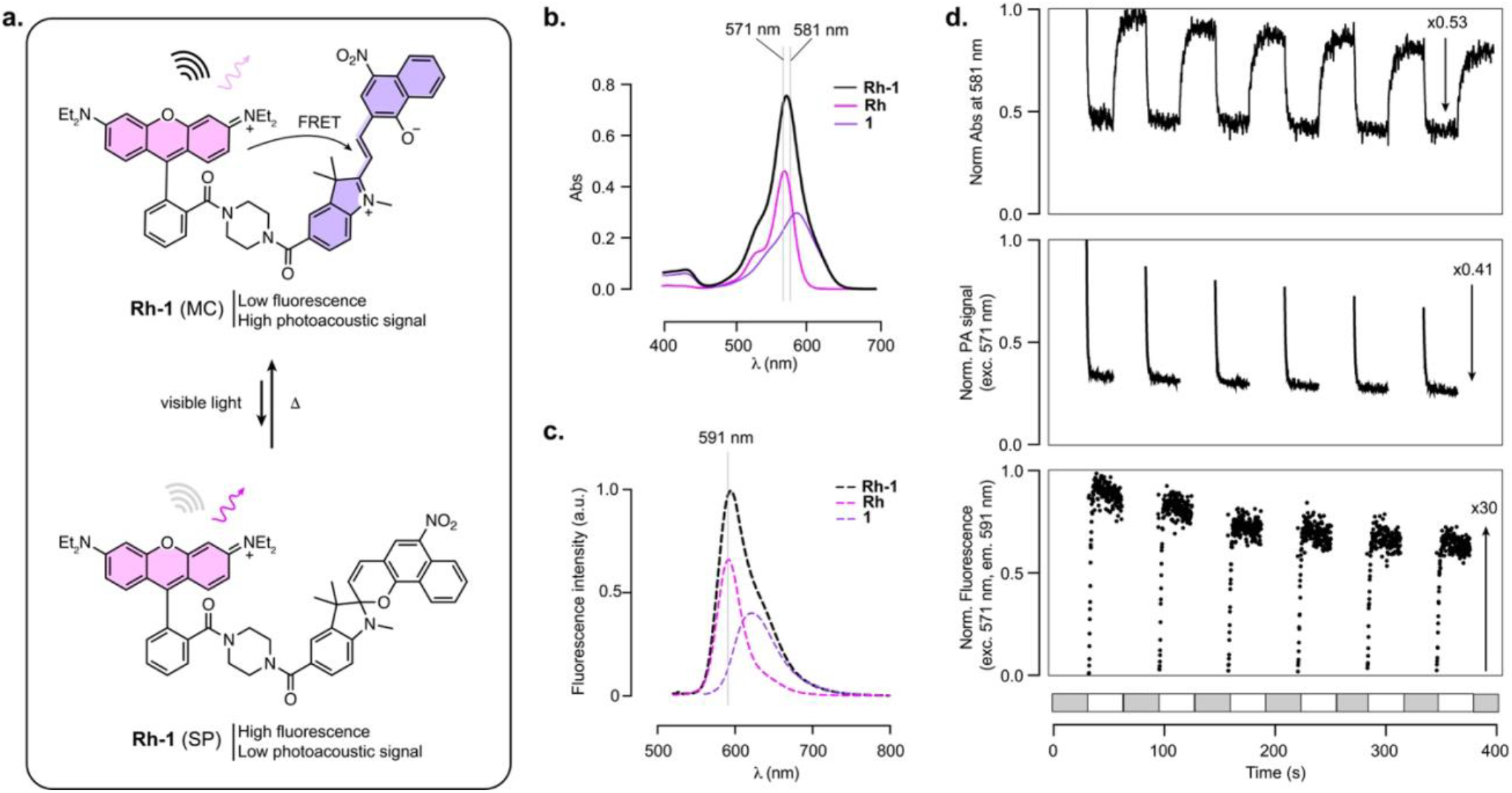
**a**. Structure and photoisomerization of dyad **Rh-1. b**. Absorption spectrum of dyad **Rh-1**, and parent building blocks **Rh** and **1** (10 μM, DMSO). **c**. Normalized fluorescence spectrum of **Rh-1** and deconvoluted contributions of the fluorescence of **Rh** and **1** to the full spectrum. **d**. Normalized absorbance, photoacoustic signal and fluorescence signal of **Rh-1** upon cycles of illumination and dark. Grey and white boxes underneath correspond to dark and illumination timeframes, respectively. Illumination: 30s, dark: 30s.

### 3. CONCLUSION

In this work, we introduce spironaphthopyrans as a new family of photoswitches for multimodal imaging. We synthesized and characterized a small library of substituted switches and systematically investigated their photoswitching properties. We highlight key difference in SP/MC equilibrium, switching amplitude and kinetics, and identify nitro-spironaphtopyran **1** as a robust negative photoswitch. Incorporation of this compound into a rhodamine–spiropyran dyad yielded a FRET-based probe that toggles reversibly between fluorescence- and photoacoustic-dominant states, providing, to the best of our knowledge, the first demonstration of SP/MC-based probes characterized for photoacoustic imaging. This study highlights the potential of spironaphthopyrans as versatile optical switches for dynamic, dual-modality contrast, yielding a probe with rapid switching and large dynamic range. Importantly, the high contrast photoswitching in either modality alone presents interesting perspectives for this compound.[4, 39] While further refinement of switching kinetics and photostability will be required for biological applications, we envision that strategies such as nanoparticle embedding[40, 41] or integration with self-labeling protein tags[42] will offer opportunities to adapt them to advanced imaging techniques towards multimodal fluorescence/photoacoustic imaging.

## SUPPLEMENTARY MATERIAL

Supplementary figures and tables, methods, synthesis and characterization for all new compounds (pdf).

## ACKOWLEDGMENTS

This work was supported by the European Molecular Biology Laboratory (EMBL) and the Chan Zuckerberg Initiative (Deep Tissue Imaging grants no. 2020-225346 and no. 2024-337799). We acknowledge the support of the EMBL Metabolomics Core Facility (MCF) for the acquisition and analysis of liquid chromatography-mass spectrometry data, and the EMBL Mechanical and Electronic Workshops for parts. We thank Dr. Richard Lincoln (EMBL) for critical reading of this manuscript, and Dr. Thomas Quail (EMBL) for contributive discussions.

## ABBREVIATIONS

Abs: absorption
DMSO: dimethyl sulfoxide
FL: fluorescence
FRET: Förster Resonance Energy Transfer
MC: merocyanine
PA: photoacoustic
SP: spiropyran.

