## Supporting Information for "Bridging Light and Sound: a Spironaphtopyran-Rhodamine Dyad with High-Contrast Photoswitching Between Fluorescence and Photoacoustic Signal"

#### Supplementary Figures and Tables

**Figure S1.** Synthesis of compounds **1-5** from substituted naphtols. Percentages indicate yields of the isolated purified compounds.

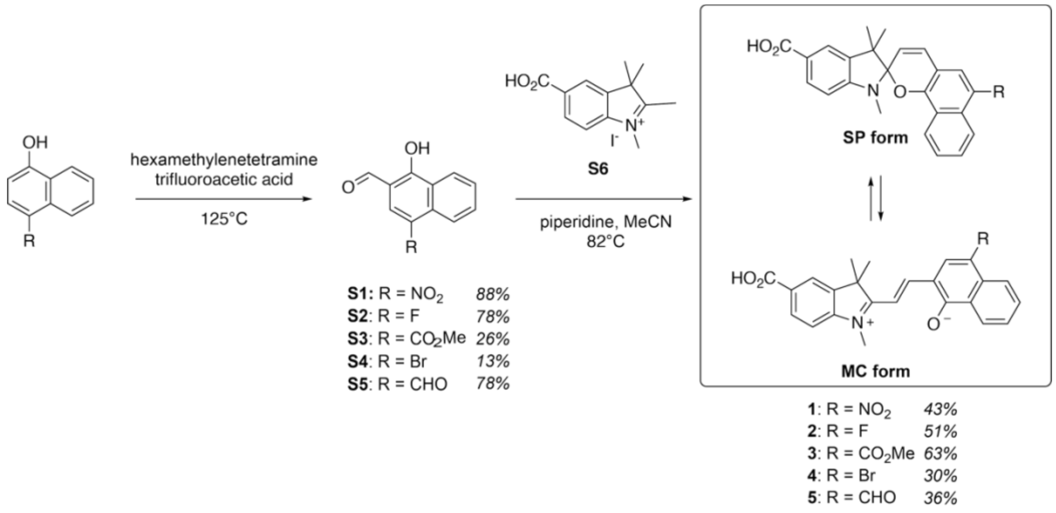

**Figure S2.** Absorption spectra of compounds **1-5** (10 μM) in different solvents.

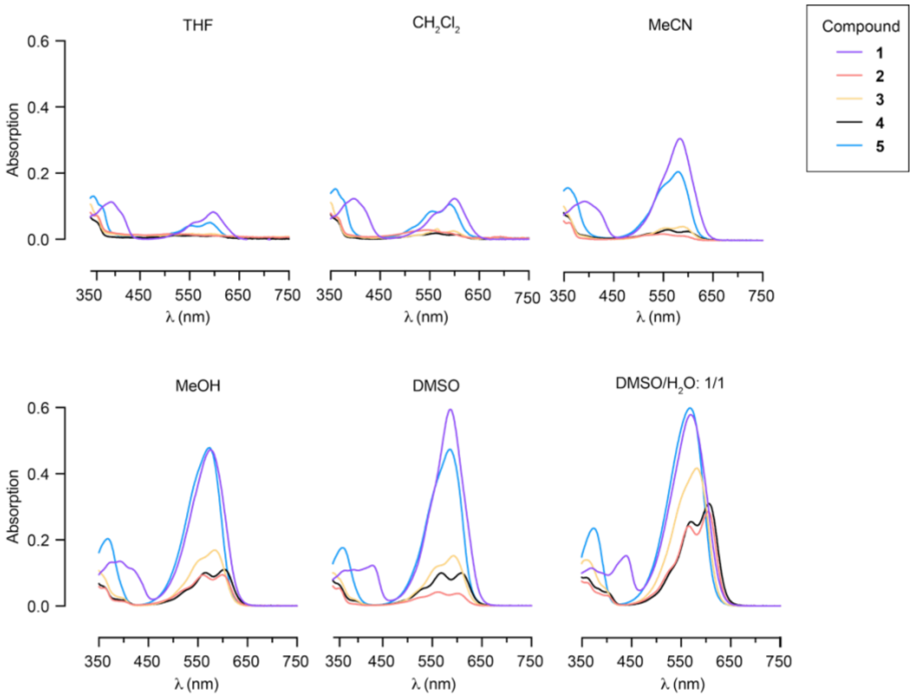

**Figure S3**. Normalized absorption and fluorescence spectra of compounds **1-5** in anhydrous DMSO.

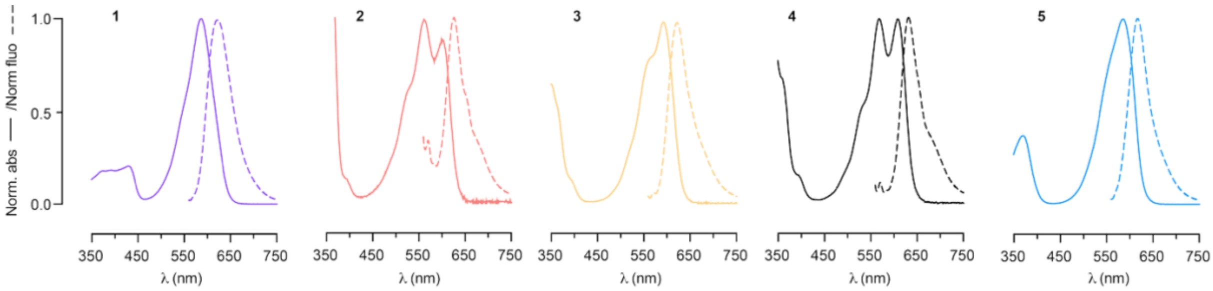

**Figure S4.** Determination of the SP/MC ratio of compounds **1-5** in the dark state by $^{1}H$ NMR in anhydrous $(CD_3)_2SO$. The peaks used for integration are annotated in each spectrum. The peaks corresponding to the gem-dimethyl group were used to determine the major isomer, and well-separated peaks in the aromatic region (integrated in the cropped NMR spectra below) were additionally used to calculate the ratio between the two species.

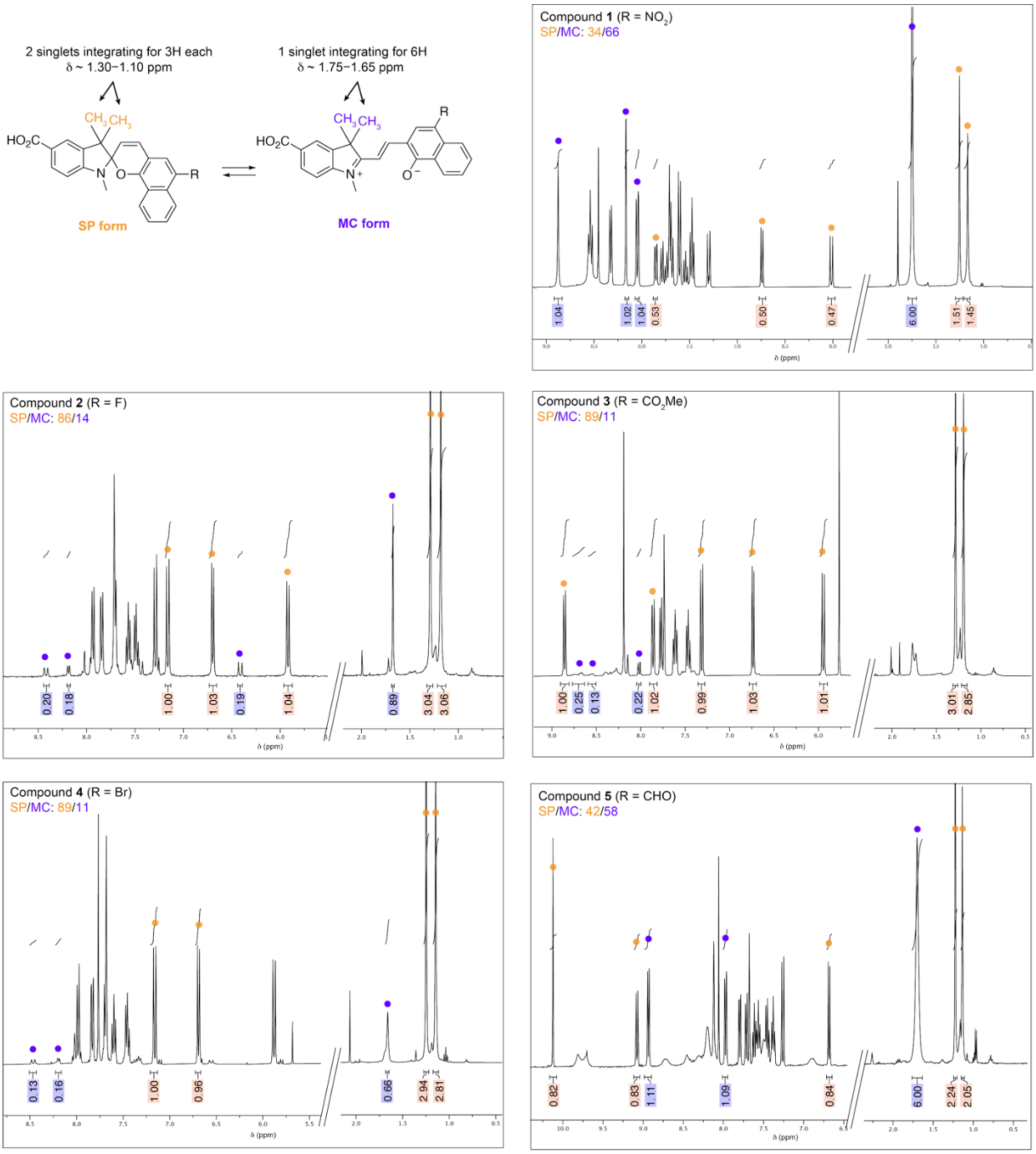

**Figure S5. a**. Aqueous stability of compounds **1-6** in $H_2O$.

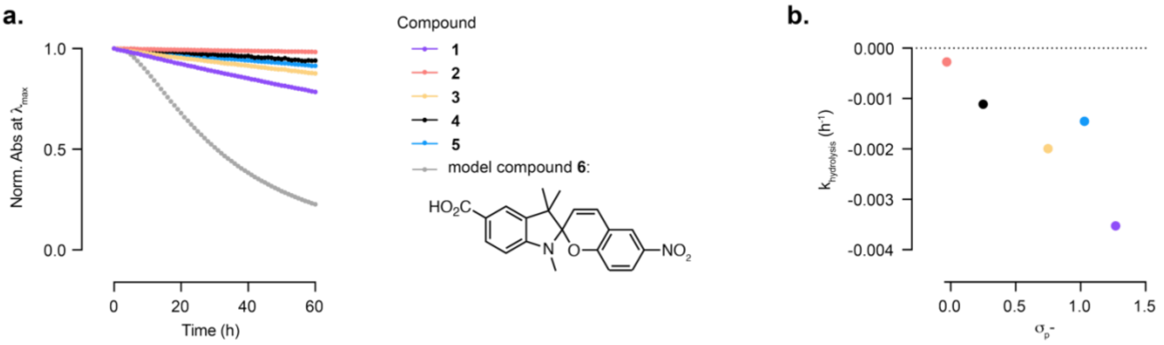

**Figure S6**. Schematic (**a**.) and picture (**b**.) of the multimodal set-up built for photoswitching characterization in Abs/FL/PA.

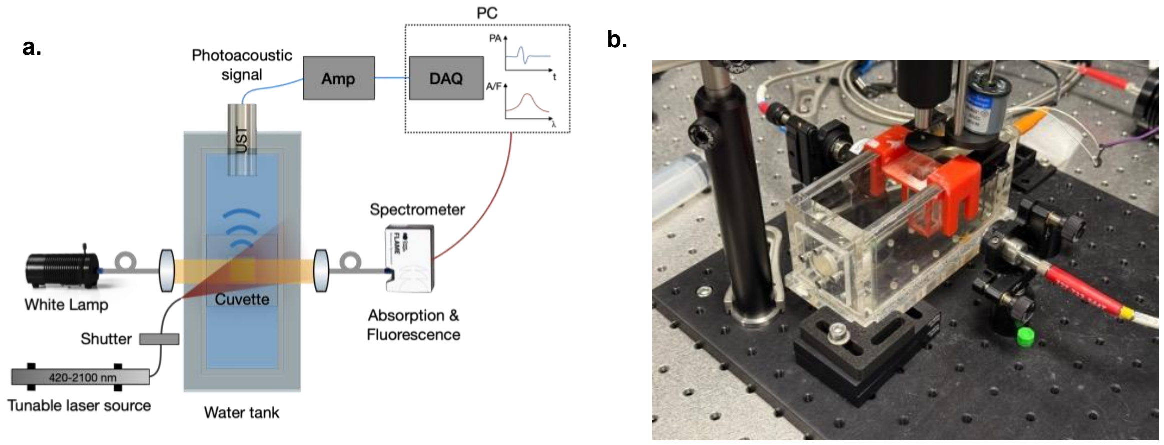

**Figure S7**. Absorbance of compound **5** upon cycles of illumination and dark with insert corresponding to a zoom-in on the area of exchange in photoswitching directionality. Grey and white boxes underneath correspond to dark and illumination frames, respectively. Illumination: 60s, dark: 60s.

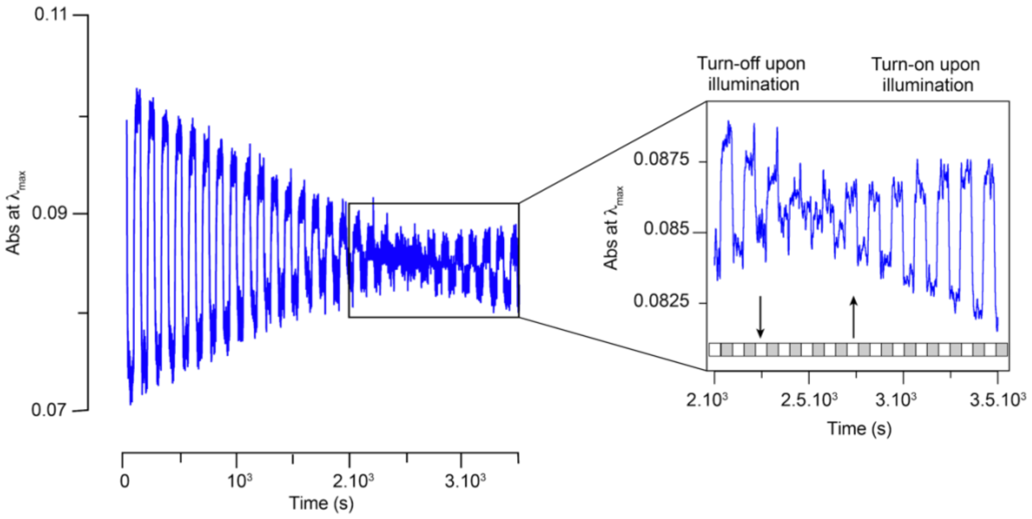

**Figure S8.** Proposed simplified photoisomerization scheme involving formation of a metastable cis-MC isomer absorbing in the visible, which thermally relaxes to the trans-MC.

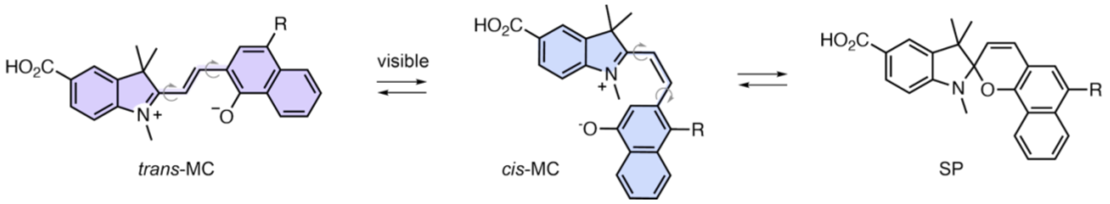

**Figure S9.** Representative turn-on and turn-off profiles for compounds **1-5** (**a.-e**.) with mono-exponential fit used to obtain kinetics parameters. The full photoswitching curves can be found in Figure 3.

**a.** Compound **1**

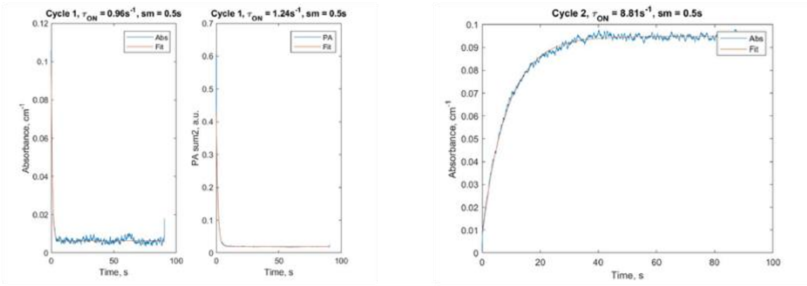

**b.** Compound **2**

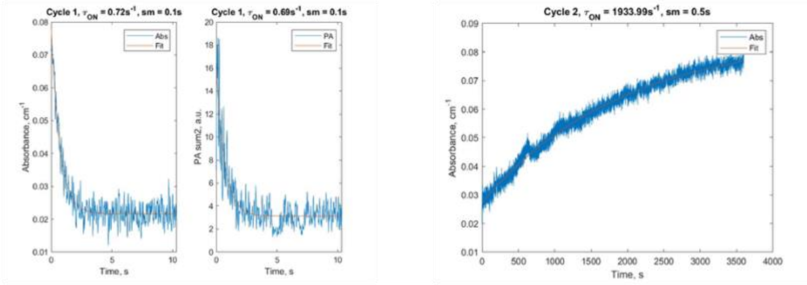

**c.** Compound **3**

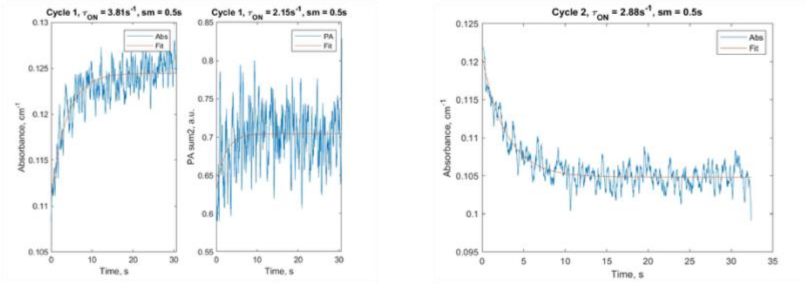

**d.** Compound **4**

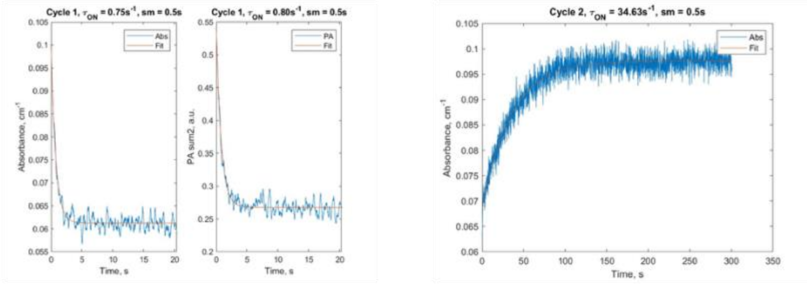

**e.** Compound **5**

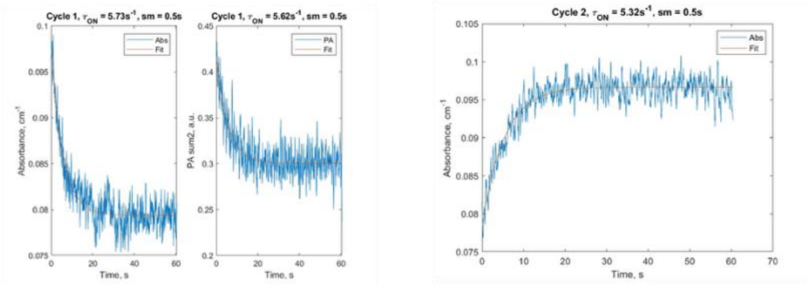

**Figure S10.** Effect of water addition to the photoisomerization and thermal relaxation kinetics (**a**.) and the dynamic range of photoswitching (**b**.) for compound **2**.

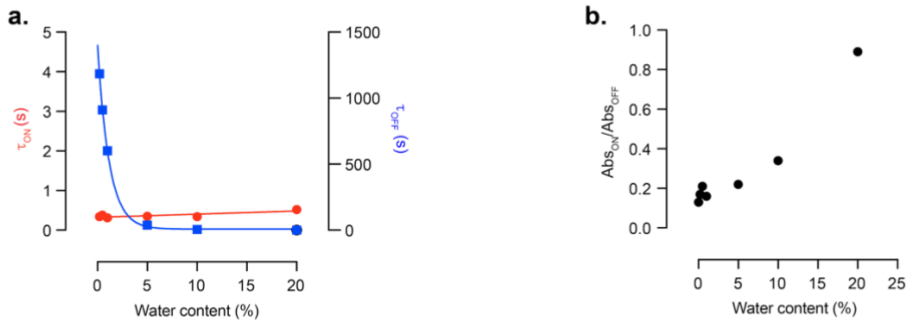

**Figure S11. a.** Structure of compounds **1** and **Rh**. **b**. Absorption spectra of compounds **1** and **Rh** and normalized fluorescence spectrum of **Rh** showing spectral overlap.

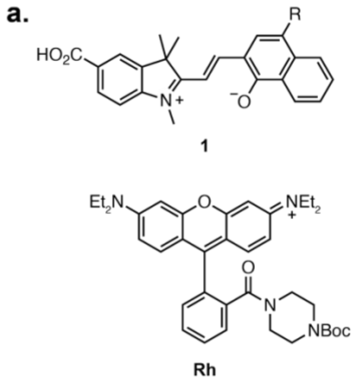

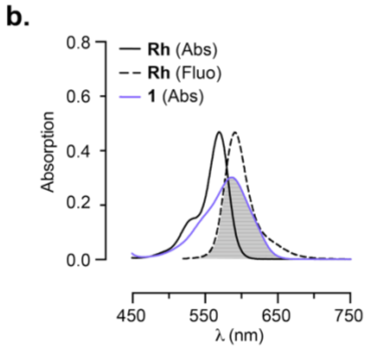

**Figure S12.** Synthesis of photoswitchable dyad **Rh-1**.

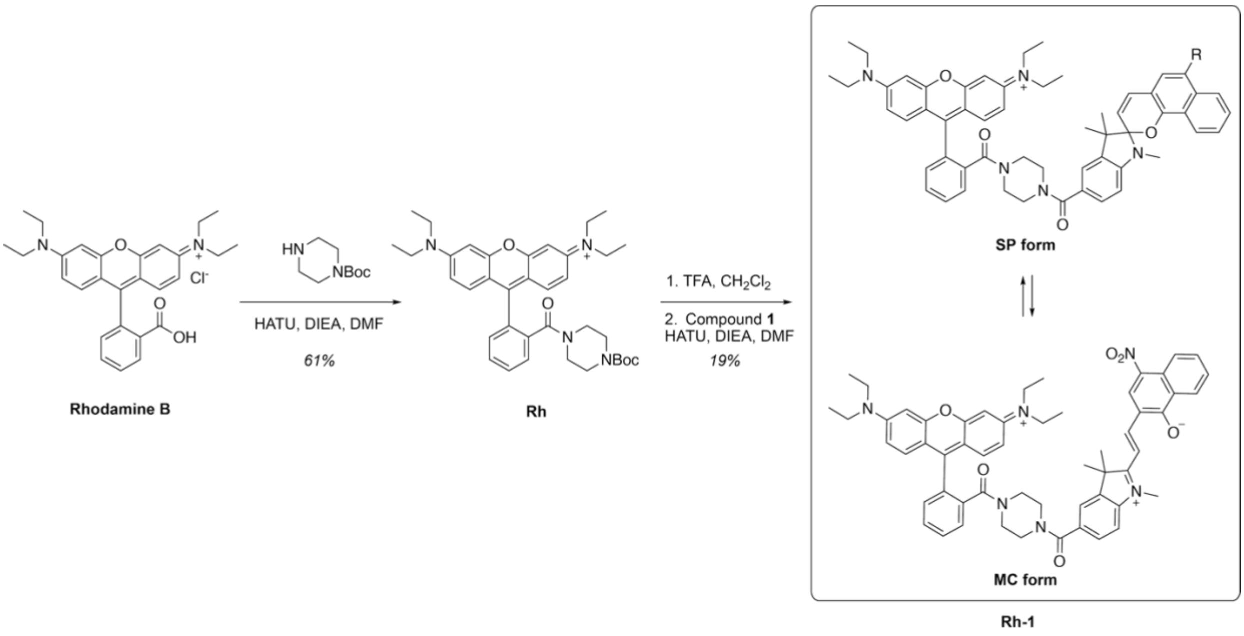

**Figure S13**. Determination of the SP/MC ratio of compound **Rh-1** in the dark state by $^{1}H$ NMR in anhydrous $(CD_3)_2SO$. The peaks used for integration are annotated in the spectrum.

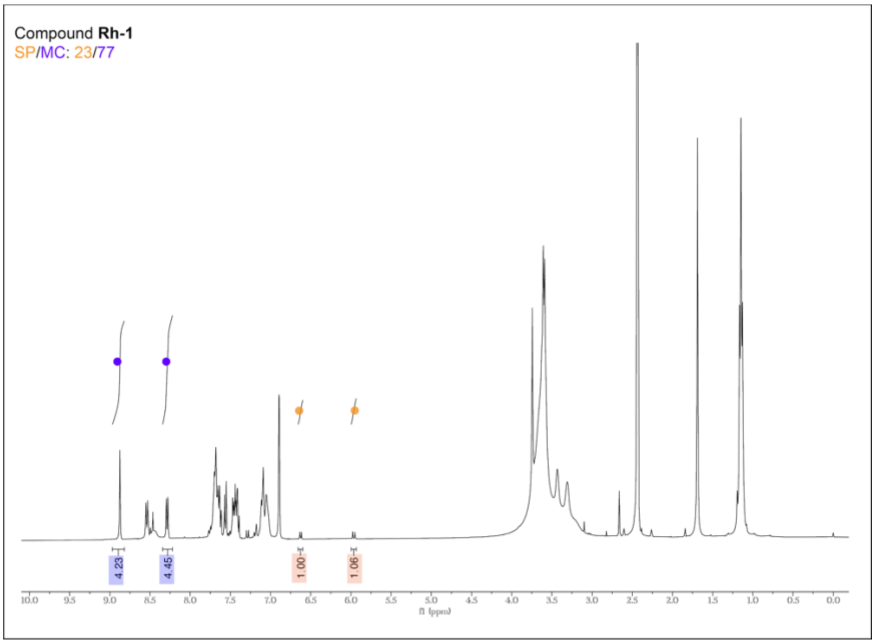

**Figure S14.** Representative photoactivation (**a**.) and thermal relaxation (**b**.) profiles for compound **Rh-1** with mono-exponential fit used to extract kinetics parameters.

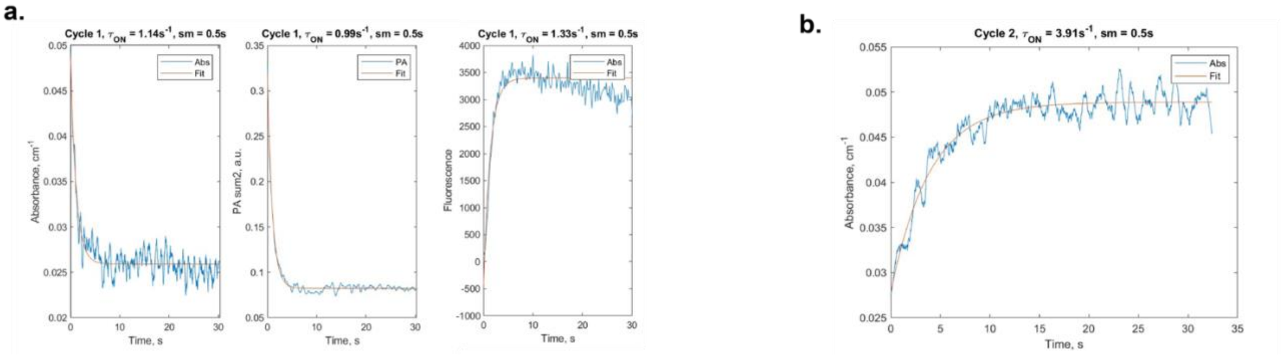

**Figure S15.** Normalized photoacoustic signal and fluorescence signal of **Rh** upon cycles of illumination and dark. Grey and white boxes underneath correspond to dark and illumination timeframes, respectively. Illumination: 30s, dark: 30s.

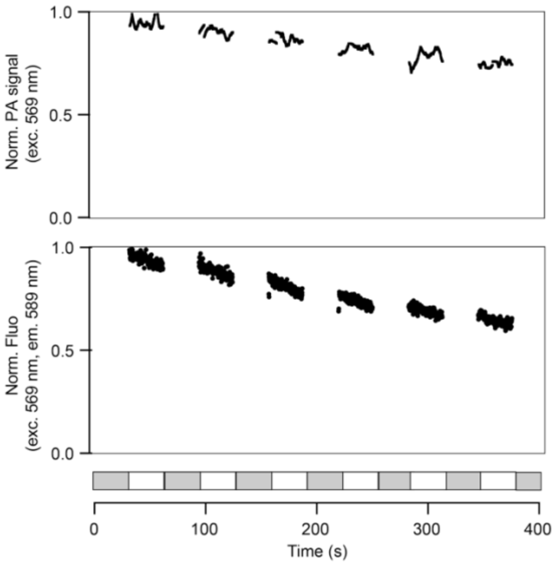

**Figure S16.** Fluorescence intensity of compounds **Rh** (**a**.) and **Rh-1** (**b**.) during cycle of illumination/dark (30s/30s). Curves were fitted to a mono-exponential to obtain photobleaching rates (**c**.).

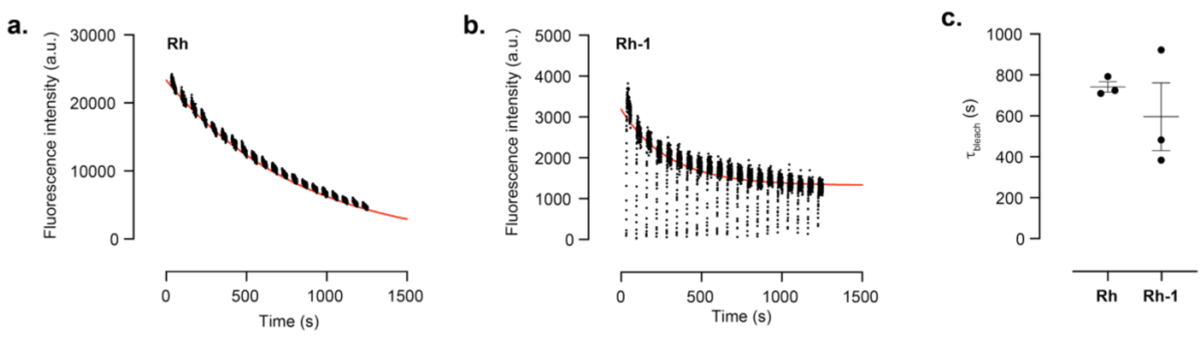

**Figure S17.** Effect of water addition to the photoisomerization and thermal relaxation kinetics (**a.**) and the dynamic range in absorbance (**b.**) for compound **Rh-1**. Mean ±SEM.

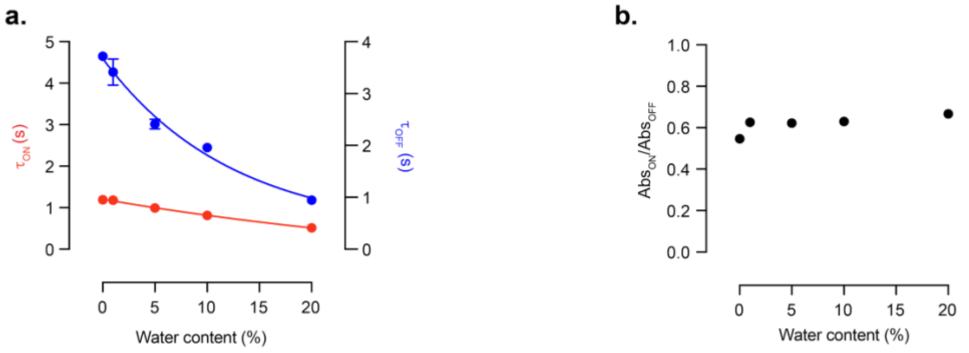

#### Supplementary Note: Molecular modelling

Density functional theory (DFT) and Ttme-dependent density functional theory (TDDFT) were performed with with Orca 6.0,[1] using the B3LYP functional and 6-31G(d,p) basis set, including solvent effects via the CPCM continuum model. Geometry optimizations and energy calculations were carried out in DMSO to identify the most stable isomers in solution. Merocyanines can adopt either planar, trans or bent, cis configurations, with the trans isomers being more stable (Figure S18).[2] For compound **1**, the TTC and TTT trans isomers were the most stable, whereas the cis forms were substantially destabilized ($\Delta E$ = 8.9−13.9 kcal·mol$^{-1}$, with the TCC form collapsing to SP). As observed experimentally, the SP form itself was less stable, about 5 kcal·mol$^{-1}$ higher in energy (Figure S19), consistent with significant MC stabilization in DMSO. In contrast, analogous calculations for compounds **2** and **3** yielded smaller MC-SP energy gaps ($\Delta E$ = 3.2 and 3.5 kcal·mol$^{-1}$, respectively, Figure S20), in line with their larger shift towards the SP form. TDDFT on the optimized geometries of compound **1** (TTC and TTT) revealed that the absorption band in the visible region of the spectrum arises from a HOMO→LUMO, $\pi\rightarrow\pi^*$ transition (Figure S21). The computed absorption maxima for TTC and TTT differ by ~20 nm, supporting that the shoulders observed on the experimental absorption spectra could be attributed to the coexistence of multiple MC isomers. However, the calculated transition energy (512 nm) is significantly blue-shifted relative to the experimental value (586 nm), and using different functional and basis set did not reduce this discrepancy. Given the pronounced environmental sensitivity of these compounds, we hypothesize that this deviation might be due to the implicit solvent model used in these calculations, which might not fully recapitulate solvent effects here.

**Figure S18**. Molecular structures of the different merocyanine and spiropyran isomers of compounds **1-5**. The isomer notation describes the cis or trans configuration of bonds $\alpha$, $\beta$, $\gamma$. E.g. TTC describes the isomer with $\alpha$ in trans, $\beta$ in trans and $\gamma$ in cis.

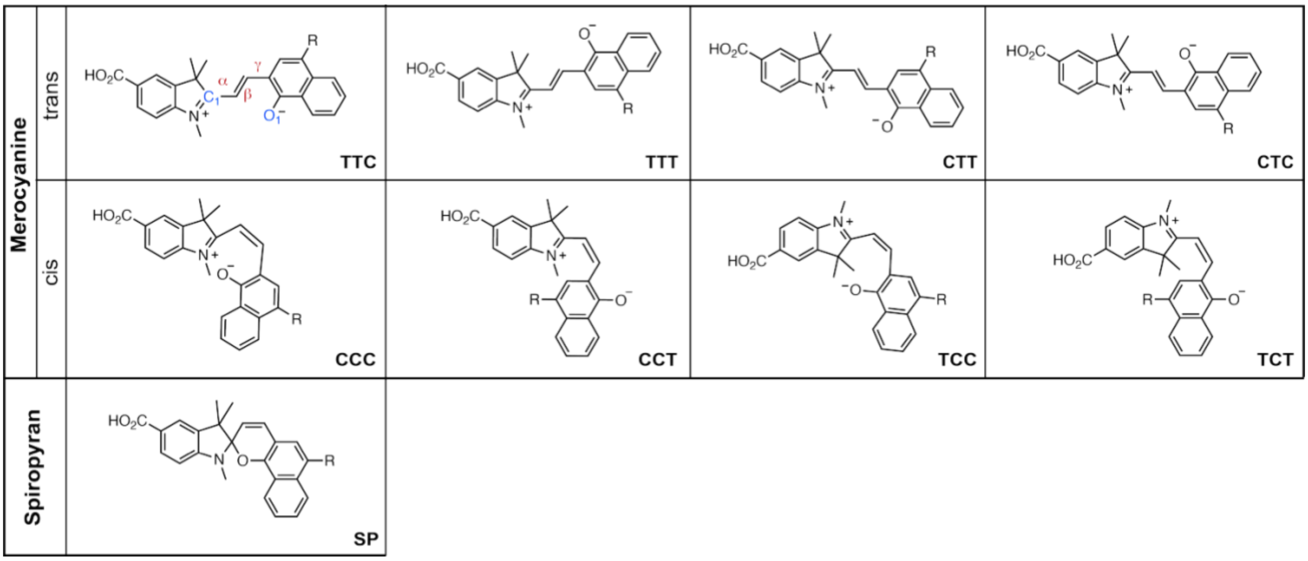

**Figure S19**. **a**. Properties of the calculated molecular structures of the isomers of compound   in DMSO: dihedral angles $\alpha$, $\beta$, $\gamma$, distance between C1 and O1 (see Figure S6 for numbering), and relative energies. [a]Electronic energy differences and [b]Gibbs free energy differences at 298K, compared to the most stable isomer TTC. **b**. Calculated molecular structures (B3LYP, 6-31G(d,p) of the isomers of compound **1** in DMSO.

**a.**

| | Isomer | α (°) | β (°) | γ (°) | $C_1$-$O_1$ (Å) | ΔE (kcal·mol$^{-1}$)[a] | ΔG (kcal·mol$^{-1}$)[b] |
|---|---|---|---|---|---|---|---|
| trans | TTC | 179.5 | 179.6 | 0.3 | 4.2 | 0 | 0 |
| trans | TTT | -179.7 | -179.8 | 179.9 | 5.0 | 0.5 | 0.4 |
| trans | CTT | -9.0 | -179.6 | 178.4 | 5.1 | 2.1 | 2.2 |
| trans | CTC | 14.5 | 177.7 | 2.9 | 4.2 | 1.8 | 2.0 |
| cis | CCC | -37.2 | -17.8 | -16.2 | 2.8 | 8.9 | 8.8 |
| cis | CCT | -29.7 | -21.1 | 174.1 | 5.3 | 9.6 | 8.8 |
| cis | TCC | | converges to SP | | | / | / |
| cis | TCT | -163.8 | 32.8 | -167.9 | 5.2 | 13.9 | 14.2 |
| | SP | 120.8 | -1.3 | -0.4 | 1.5 | 5.0 | 5.9 |

**b.**

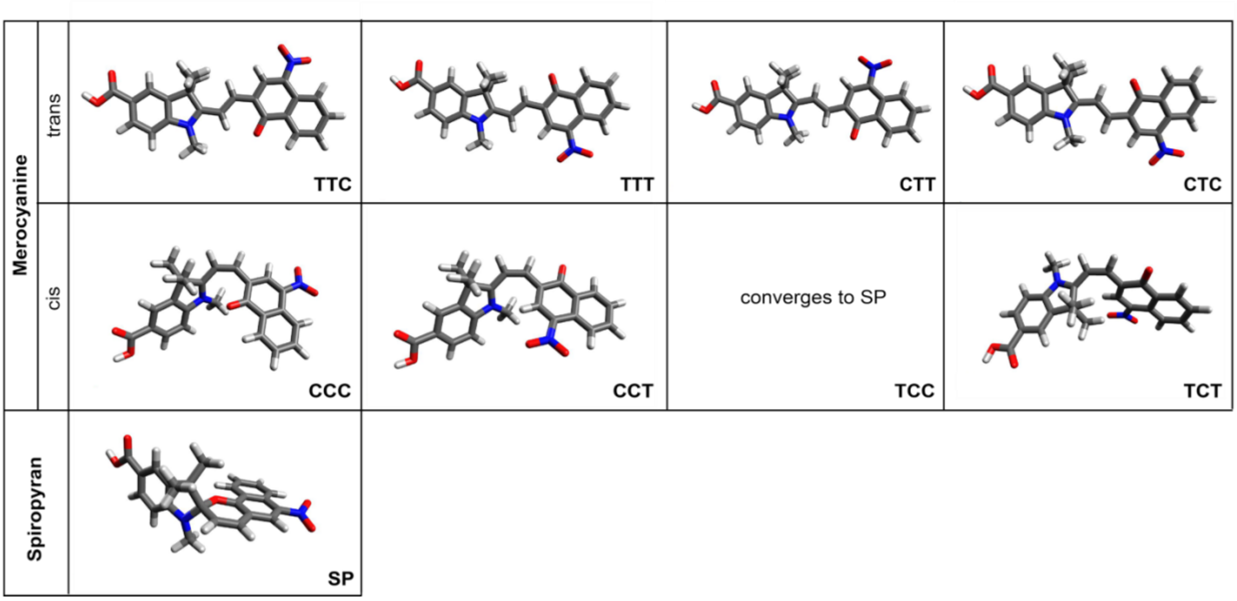

**Figure S20**. Electronic energy differences for TTC, TTT and SP isomers for compounds **1**, **2** and **3**.

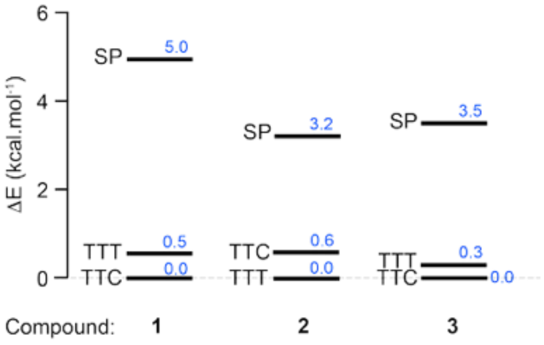

**Figure S21. a**. Calculated visible electronic transition for the TTC and TTT isomers of compound **1** in DMSO. **b**. Representation of the HOMO and LUMO of the TTC and TTT isomers of compound **1**.

**a.**

| Isomer | λ (nm) | Oscillator strength | Main transition |
|---|---|---|---|
| TTC | 494 | 1.55 | HOMO → LUMO 96% |
| TTT | 512 | 1.49 | HOMO → LUMO 96% |

**b.**

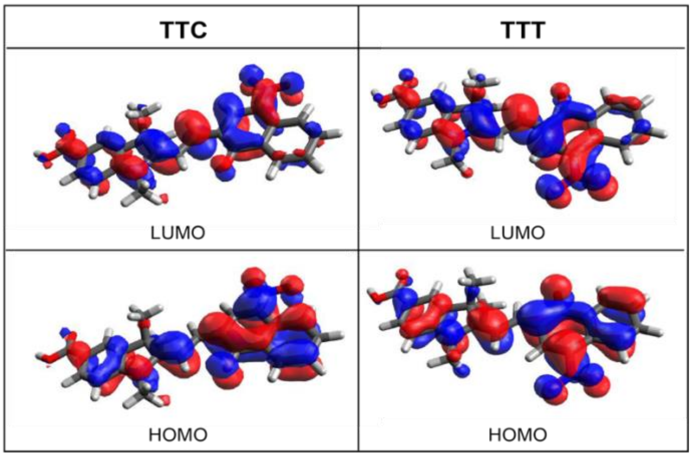

#### General Experimental Information

#### Synthesis

Commercial reagents were obtained from reputable suppliers (e.g. Merck, TCI, BLDpharm) and used as received. All solvents used for chemical reactions were of anhydrous grade, purchased in septum-sealed bottles stored under an inert atmosphere. Reactions under an inert atmosphere were sealed with septa and purged under vacuum/argon on a Schlenk line. Reactions were performed in round-bottomed flasks or septum-sealed vials.

Reactions were monitored by thin layer chromatography (TLC) on precoated aluminium plates (silica gel 60 F254, 200 µm thickness) or by LCMS (Agilent 1260 Infinity II; ZORBAX SB-C18 18 µm 80 Å, 2.1x50 mm column, 5 to 20 µL injection, 5–95% $CH_3CN/H_2O$ gradient with constant 0.1% v/v $HCO_2H$ additive, 8 to 10 min run, 0.6 mL/min flow, ESI positive ion mode, detection at 254 nm). TLC plates were visualized by UV illumination (254 nm). Compounds were purified by flash chromatography on an automated purification system (Biotage Isolera One) using pre-packed silica cartridges (Biotage Sfär Duo, 60 Å pores, 60 µm particles size) or by preparative HPLC (Agilent 1260 Infinity II, Phenomenex Gemini NX 21.2x150 mm, 5 µm C18 column). High-resolution mass spectrometry was performed by the EMBL Metabolomics Core Facility.

NMR spectra were recorded on a 400 MHz spectrometer (Bruker 400 UltraShield) in deuterated solvents. All spectra were recorded at 298 K. $^1H$ and $^{13}C$ chemical shifts (δ) were referenced to residual solvent peaks. Data for $^1H$ NMR spectra are reported as chemical shift (δ in ppm), multiplicity (s = singlet, d = doublet, t = triplet, q = quartet, p = pentuplet, dd = doublet of doublets, ddd= doublet of doublets of doublets, dt = doublet of triplets, tt = triplet of triplets, dtt = doublet of triplets of triplets, m = multiplet, br s = broad signal), coupling constant (*J* in Hz), integration. Data for $^{13}C$ NMR spectra are reported by chemical shift (δ in ppm). Data was processed using Mnova from Mestrelab.
*Note*: In the case of spiropyran/merocyanine compounds **1**-**5**, the deuterated solvent for NMR measurement was carefully chosen to shift the equilibrium strongly towards the SP form, to facilitate analysis of the NMR spectrum with a single isomer in solution and provide clearer evidence of identity and purity. In several cases, this required highly apolar solvents ($CDCl_3$, dioxane-d4) in which these $CO_2H$-substituted compound display low solubility. As such, $^{13}C$ NMR is not reported for compounds **1**-**5** due to issues of solubility, and we report $^1H$ NMR, HPLC trace and HRMS ad evidence of molecular identify and purity.

#### UV-Vis, Fluorescence and Photoacoustic Spectroscopy

The following notations are used:
Abs: absorption
Fl: fluorescence signal
PA: photoacoustic signal
OFF state: dark state
ON state: PSS obtained following illumination at $\lambda_{max}$
$\lambda_{max}$ (nm): wavelength at maximal absorption
$\lambda_{em}$ (nm): wavelength at maximal fluorescence emission
ε ($M^{-1}\cdot cm^{-1}$): molar extinction coefficient.
$\Phi_{Fl}$: fluorescence quantum yield
$\tau_{ON}$: time constant corresponding to a mono-exponential fit of the photoactivation
$\tau_{OFF}$: time constant corresponding to a mono-exponential fit of the thermal relaxation

###### Characterization of the dark state of the probes

All measurements were performed at room temperature (23 ± 2°C). Compounds were prepared as stock solutions at 2 mM in DMSO which were diluted in solvents. All organic solvents used were anhydrous of spectroscopy grade. Spectroscopy in organic solvents was performed in 1-cm path length quartz cuvettes were used (Hellma Suprasil Quartz).

Absorption spectra were recorded on a Cary Model 60 spectrophotometer (Agilent). Fluorescence spectra were recorded on a JASCO spectrofluorometer (FP-8500). Data was analysed and graphs were plotted using Prism (GraphPad). Absorption spectra, maximum absorption wavelength ($\lambda_{max}$), extinction coefficient (ε) and maximum emission wavelength ($\lambda_{em}$) were measured in triplicate under the following conditions, reported values for ε are the mean of 3 replicates. Spectra were recorded at 10 µM concentration for absorption spectra measurements, and diluted so that A<0.1 for fluorescence spectra measurements.

Fluorescence quantum yields ($\Phi_F$) were measured using an absolute quantum yield measurement system (Quantaurus, C11347, Hamamatsu). Measurements were carried out using dilute samples (A<0.1), self-absorption corrections were performed using the Quantaurus software.

**Characterization of the photoswitching behavior of the probes with a multimodal spectroscopy setup.**

A custom-built multimodal spectroscopy system was used to simultaneously measure photoacoustic (PA), absorbance (Abs), and fluorescence (Fl) signals from samples in solution (schematic in Figure S10). The setup was an extension on our previous system,[3] and consisted of a water tank with a cuvette holder, a tunable excitation laser with a computer-controlled shutter, a focused ultrasound transducer, a broadband white-light source, and a spectrometer. In the present configuration with a single spectrometer, the system was operated either in PA + Abs mode or in PA + Fl mode.

*Photoacoustic measurements.*

PA signals were excited at $\lambda_{max}$ using a 100 Hz pulsed tunable OPO laser (EVO I, Innolas Laser GmbH) directed onto the cuvette from above. A custom shutter provided precise control over illumination timing. Acoustic waves generated in the sample were detected with a focused 0.5 MHz water-immersion transducer (V301, Olympus IMS), amplified (DHPVA-101, FEMTO Messtechnik GmbH), and digitized with a 125 MSa/s DAQ card (ATS9440, Alazar Technologies Inc.).

*Fluorescence measurements.*

Fluorescence excitation was provided by the same OPO laser source at $\lambda_{max}$. Emission was collected with a multimode fiber–collimator assembly and directed into a spectrometer (Flame-T-VISNIR, Ocean Insight Inc.) operated in triggered mode and the trace was recorded at $\lambda_{em}$. Due to low numerical aperture collection and relatively low sample concentrations, the integration time was set to 30–100 ms, corresponding to the signal integrated over 3–10 excitation pulses.

*Absorbance measurements.*

Absorbance was measured by monitoring the transmission of a broadband white-light source (HL2000-HP, Ocean Insight Inc., 360–1700 nm) through the cuvette using the same spectrometer. The sample was excited at $\lambda_{max}$ and the absorbtion trace was recorded at $\lambda_{max}$ plus 10 nm shift to avoid overlap of the excitation pulse. A dark spectrum was first recorded to account for detector noise, followed by a reference spectrum of the buffer. Absorbance spectra were then calculated as $Abs(\lambda) = -\log_{10}(I_{sample}(\lambda) - I_{dark}(\lambda))/(I_{ref}(\lambda) - I_{dark}(\lambda))$. To avoid overlap between fluorescence and absorbance signals, the spectrometer was triggered with a 1 ms delay relative to the excitation pulse, and its exposure time was fixed to 5 ms.

**Data processing**

A MATLAB-based pipeline was developed to jointly analyze Abs/FL and PA traces. The workflow included segmentation of illumination cycles, normalization of PA signals, extraction of scalar PA metrics, and kinetic fitting of ON/OFF segments.

***Segmentation of absorbance/fluorescence traces.***

Trace files contained time stamps, intensity values, and a binary illumination flag (1 = ON, 0 = OFF, corresponding to shutter state). Transitions in the flag were used to segment the data into ON and OFF intervals. Each segment was stored with its raw trace, a relative time axis, and a smoothed version (Savitzky–Golay filter, 50-point window).

***Photoacoustic and laser energy data.***

For each ON segment, PA waveforms and the OPO energy log were loaded. Because PA amplitudes scale linearly with excitation energy, each PA trace was normalized on a per-pulse basis by the interpolated energy log.

***Kinetic parameters extraction.***

A temporal window corresponding to the acoustic propagation through the cuvette (estimated from the speed of sound and cuvette geometry) was selected. The squared PA amplitude within this window was summed for each pulse, yielding a scalar PA energy metric. Both raw and normalized metrics were calculated and interpolated onto the Abs/FL time axis for direct comparison.

***Fitting procedures.***

***Abs + PA (Photoswitches 1–5):*** ON-segment for Abs and PA traces were fitted with a mono-exponential function, and OFF-segment represented only by Abs measurement were also fitted mono-exponentially

$$Abs(t) = Abs_0 + \Delta Abs e^{-t/\tau_{ON/OFF}};\ PA(t) = PA_0 + \Delta PA e^{-t/\tau_{ON}}$$

***Abs + PA (Rh, Rh-1):*** As bleaching effects were negligible, absorbance traces were directly fitted with a mono-exponential (parameters $Abs_0$, $\Delta Abs$ and $\tau_{ON/OFF}$).

***FL + PA (Dyad)*:** Because fluorescence bleaching was significant even within a single cycle, the long-term fluorescence decay was first fitted globally with an exponential bleaching model.

$$FL_b(t) = FL_0 + \Delta FL_b e^{-t/\tau_b}$$

Fluorescence traces within each ON segment were then corrected by this bleaching trend $F_b$, then fitted with a mono-exponential $FL(t) = FL_0 + \Delta FL e^{-t/\tau_{ON}}$ to extract $FL_0$, $\Delta FL$ and $\tau_{ON}$. PA metrics were fitted analogously (mono-exponential, yielding $PA_0$, $\Delta PA$ and $\tau_{ON}$). No PA or fluorescence signals were acquired during OFF states.

#### Molecular Modelling

Density functional theory (DFT) calculations were performed with Orca 6.0[1] at the B3LYP/6-31G(d,p) level. The effect of solvent (DMSO or THF) was accounted by continuum CPCM for geometry optimization, energy and frequency calculations. True minima were confirmed by the absence of imaginary frequencies for all calculated structures. Electronic transitions were obtained by time-dependent density functional theory (TDDFT) method implemented in Orca 6.0, using DMSO as solvent.

Optimized molecular structures and orbitals were visualized using Avogadro 1.2.1.[4]

#### FRET Efficiency Calculation

The FRET efficiency (E) between donor **Rh** and acceptor **1** was calculated using the following equation[5]:

$$E = \frac{1}{1 + (\frac{r}{R_0})^6}$$

Where: $r$ is the distance between donor and acceptor; $R_0$ is the Förster radius corresponding to a distance at which the energy transfer efficiency is 50%.

The Förster radius $R_0$ was calculated as follows:

$$R_0 = 0.211[\kappa^2 \Phi_{FL} n^{-4} J]^{1/6} \quad \text{in Å}$$

$$J(\lambda) = \int_0^\infty FL_D(\lambda)\varepsilon_A(\lambda)\lambda^4 d\lambda \Big/ \int_0^\infty FL_D(\lambda)d\lambda \quad \text{in } M^{-1}\cdot cm^{-1}\cdot nm^4$$

Where: $\kappa$ is the orientation factor; $\Phi_{FL}$ is the fluorescence quantum yield of the donor **Rh** ($\Phi_{FL}$ = 0.55); n is the refractive index of the medium (DMSO, $n$ = 1.479); $J(\lambda)$ is the overlap integral; $FL_D(\lambda)$ is the fluorescence spectrum of the donor **Rh**; $\varepsilon_A(\lambda)$ is the extinction coefficient spectrum of acceptor **1** in $M^{-1}\cdot cm^{-1}$; $\lambda$ is the wavelength in nm.

For calculation of the overlap integral, the extrapolated spectrum of the pure MC form of **1** was used. This was calculated using the absorption spectrum of **1** in the dark state and corrected by the SP/MC ratio in the dark state (34/66 here).

Using our experimental data, we obtain:

$J(\lambda)$ = 8.11.$10^{15}$ $M^{-1}.cm^{-1}.nm^4$

$R_0$ = 61.6 Å

Geometry optimization of the structure of compound **Rh-1** using B3LYP/6-31G(d) with Orca 6.0[1] led to 10 Å < $r$ < 16 Å, measured between the geometric centers of the two chromophores.

Together, this gives a FRET efficiency $E$ > 99.9% in the MC isomer of **Rh-1**.

#### Experimental and Characterizations for New Compounds

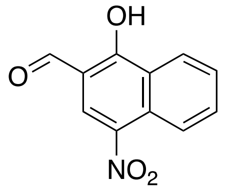

**4-nitro-1-hydroxy-2-naphthaldehyde (S1):** synthesized according to published procedure.[6]

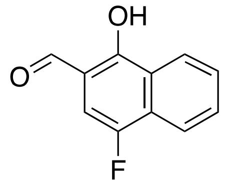

**4-fluoro-1-hydroxy-2-naphthaldehyde (S2):** 4-fluoronaphthalen-1-ol (508 mg, 3.13 mmol, 1.0 eq) and hexamethylenetetramine (440 mg, 3.14 mmol, 1.0 eq) were dissolved in trifluoroacetic acid (5.0 mL) and the mixture was heated at 126°C for 17 hours. After cooling to room temperature, the mixture was poured onto an aqueous solution of HCl 4 M (100 mL). The resulting mixture was extracted three times with $CH_2Cl_2$ and the combined organic layers were washed with an aqueous solution of HCl 4 M, brine and $H_2O$. The organic layer was dried over anhydrous $Na_2SO_4$, filtered, and evaporated to dryness to afford the title compound as a yellow solid (463 mg, 78%). $^1$H NMR (400 MHz, $CDCl_3$) δ 12.41 (s, 1H), 9.78 (s, 1H), 8.4 (d, $J$ = 8.4 Hz, 1H), 7.94 (d, $J$ = 8.4 Hz, 1H), 7.67 (ddd, $J$ = 8.3, 6.9, 1.3 Hz, 1H), 7.55 (ddd, $J$ = 8.2, 6.9, 1.2 Hz, 1H), 7.02 (d, $J$ = 9.9 Hz, 1H); $^{13}$C NMR (101 MHz, $CDCl_3$) δ 195.3, 158.2, 151.7 (d, $^1J_{CF}$ = 243 Hz), 130.9 (d, $^4J_{CF}$ = 2 Hz), 128.1 (d, $^2J_{CF}$ = 19 Hz), 127.13, 125.5 (d, $^3J_{CF}$ = 5 Hz), 124.5 (d, $^4J_{CF}$ = 3 Hz), 120.5 (d, $^3J_{CF}$ = 5 Hz), 112.5 (d, $^3J_{CF}$ = 7 Hz), 107.8 (d, $^2J_{CF}$ = 22 Hz); HRMS (ESI) calculated for $[M-H]^-$ 189.0357, found 189.0358.

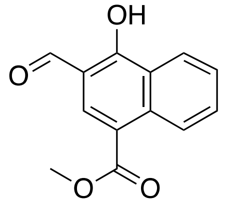

**methyl 3-formyl-4-hydroxy-1-naphthoate (S3)**: methyl 4-hydroxy-1-naphthoate (501 mg, 2.48 mmol, 1.0 eq) and hexamethylenetetramine (694 mg, 4.95 mmol, 2.0 eq) were dissolved in trifluoroacetic acid (5 mL) and the mixture was heated at 126°C for 4 days. After cooling to room temperature, the mixture was poured onto an aqueous solution of HCl 4 M (100 mL) and stirred for 2 hours. The resulting precipitate was filtered and washed with $H_2O$. The solid obtained was purified by column chromatography (0–100% EtOAc/$CH_2Cl_2$) to afford the title compound as a yellow solid (148 mg, 26%). $^1$H NMR (400 MHz, $CDCl_3$) δ 13.0 (s, 1H), 9.98 (s, 1H), 9.05 (d, $J$ = 8.7 Hz, 1H), 8.63 – 8.43 (m, 1H), 8.37 (s, 1H), 7.79 (ddd, $J$ = 8.5, 6.9, 1.4 Hz, 1H), 7.60 (ddd, $J$ = 8.2, 6.9, 1.1 Hz, 1H), 3.99 (s, 3H); $^{13}$C NMR (100 MHz, $CDCl_3$) δ 196.2, 166.8, 165.1, 135.6, 133.2, 132.5, 126.6, 126.3, 125.0, 124.7, 118.6, 113.1, 52.3; HRMS (ESI) calculated for $[M+H]^+$ 231.0652, found 231.0652.

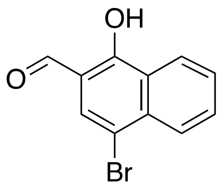

**4-bromo-1-hydroxy-2-naphthaldehyde (S4)**: 4-bromo-1-naphthol (581 mg, 2.60 mmol, 1.0 eq) and hexamethylenetetramine (368 mg, 2.63 mmol, 1.0 eq) were dissolved in trifluoroacetic acid (5 mL) and the mixture was heated at 126°C for 15 hours. After cooling to room temperature, the mixture was poured onto an aqueous solution of HCl 4 M (100 mL) and stirred for 2 hours. The mixture was extracted with $CH_2Cl_2$ (3x), and the combined organic layers were washed with HCl 4 M, brine, and $H_2O$. The organic layer was dried over anhydrous $Na_2SO_4$, filtered and evaporated. The crude product was purified by column chromatography (50–100% $CH_2Cl_2$/cyclohexane) to afford the title compound as a yellow solid (85 mg, 13%). $^1$H NMR (400 MHz, $CDCl_3$) δ 12.57 (s, 1H), 9.89 (d, *J* = 1.0 Hz, 1H), 8.45 (dd, *J* = 8.6, 1.2 Hz, 1H), 8.27 – 8.07 (m, 1H), 7.88 – 7.70 (m, 2H), 7.61 (ddd, *J* = 8.2, 7.0, 1.2 Hz, 1H); $^{13}$C NMR (100 MHz,) δ 195.3, 161.5, 135.6, 132.0, 129.6, 127.4, 127.1, 125.9, 124.9, 115.0, 112.2; HRMS (ESI) calculated for $[M-H]^-$ 248.9557, found 248.9556.

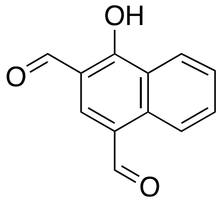

**4-hydroxynaphthalene-1,3-dicarbaldehyde (S5):** 4-hydroxy-1-naphthaldehyde (415 mg, 2.41 mmol, 1.0 eq) and hexamethylenetetramine (339 mg, 2.42 mmol, 1.0 eq) were dissolved in trifluoroacetic acid (5.0 mL) and the mixture was heated at 126°C for 18 hours. After cooling to room temperature, the mixture was poured onto a solution of HCl 4 M (100 mL) and stirred for 3 hours. The resulting precipitate was filtered and washed with $H_2O$. The solid was dried under vacuum to afford the title compound as a yellow solid (374 mg, 78%). $^1$H NMR (400 MHz, $(CD_3)_2SO$) δ 10.32 (s, 1H), 10.21 (s, 1H), 9.23 (d, *J* = 8.5 Hz, 1H), 8.54 – 8.47 (m, 1H), 8.46 (s, 1H), 7.92 (ddd, *J* = 8.4, 6.9, 1.4 Hz, 1H), 7.74 (ddd, *J* = 8.2, 6.9, 1.2 Hz, 1H); $^{13}$C NMR (100 MHz, $CDCl_3$) δ 195.8, 191.3, 166.7, 139.9, 134.5, 133.7, 127.6, 125.4, 124.86, 124.85, 124.7, 113.6; HRMS (ESI) calculated for $[M+H]^+$ 201.0546, found 201.0546.

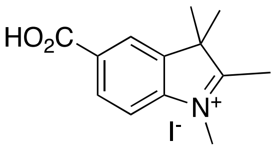

**1,2,3,3-tetramethyl-5-carboxy-3H-indolium iodide (S6):** synthesized according to published procedure.[7]

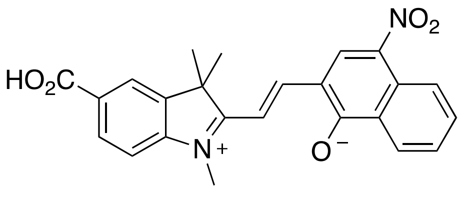

**6-nitro-1',3',3'-trimethylspiro[benzo[h]chromene-2,2'-indoline]-5'-carboxylic acid (1):** synthesized according to published procedure.[6]

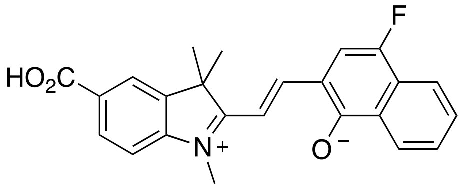

**6-fluoro-1',3',3'-trimethylspiro[benzo[h]chromene-2,2'-indoline]-5'-carboxylic acid (2):** Compound **S2** (213 mg, 1.12 mmol, 1.0 eq) and compound **S6** (387 mg, 1.12 mmol, 1.0 eq) were suspended in acetonitrile (10 mL) and piperidine (0.225 mL, 2.28 mmol, 2.0 eq) was added. The mixture was heated at 82°C for 48 hours. After cooling to room temperature, it was evaporated to dryness. The crude product was purified by column chromatography (0–100% $CH_2Cl_2$/cyclohexane containing 1% AcOH) to afford the title compound as a dark purple solid (222 mg, 51%). A small fraction of was purified by HPLC for spectroscopic measurements. $^1H$ NMR (400 MHz, $CD_3OD$) δ 7.98 – 7.87 (m, 2H), 7.81 – 7.72 (m, 2H), 7.48 (ddd, *J* = 8.3, 6.8, 1.3 Hz, 1H), 7.39 (ddd, *J* = 8.2, 6.8, 1.2 Hz, 1H), 7.08 – 7.01 (m, 2H), 6.62 (d, *J* = 8.2 Hz, 1H), 5.87 (d, *J* = 10.2 Hz, 1H), 2.80 (s, 3H), 1.36 (s, 3H), 1.23 (s, 3H); Analytical HPLC: $t_R$ = 5.00 min, purity 98%, 5–95% $CH_3CN/H_2O$, gradient with constant 0.1% formic acid additive, 8 min run, 0.6 mL/min flow, UV detection at 254 nm; HRMS (ESI) calculated for $[M+H]^+$ 390.1500, found 390.1498.

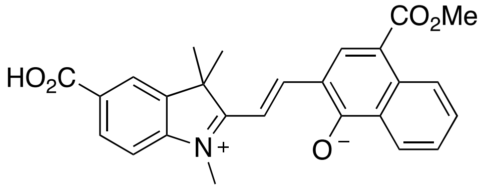

**6-(methoxycarbonyl)-1',3',3'-trimethylspiro[benzo[h]chromene-2,2'-indoline]-5'-carboxylic acid (3)**: Compound **S3** (127 mg, 0.551 mmol, 1.0 eq) and compound **S6** (191 mg, 0.552 mmol, 1.0 eq) were suspended in acetronitrile (5 mL), and piperidine (0.11 mL, 1.1 mmol, 2.0 eq) was added. The mixture was heated at 82°C for 2 days. After cooling to room temperature, it was evaporated to dryness. The crude product was purified by column chromatography (0–100% $CH_2Cl_2$/cyclohexane with 1% AcOH followed by 0–100% EtOAc/$CH_2Cl_2$ with 1% AcOH additive) to afford the title compound as a light grey solid (149 mg, 63%). $^1H$ NMR (400 MHz,$CDCl_3$) δ 8.95 (d, *J* = 8.7 Hz, 1H), 8.13 – 8.03 (m, 2H), 7.99 – 7.90 (m, 1H), 7.85 (d, *J* = 1.7 Hz, 1H), 7.55 (ddd, *J* = 8.5, 6.8, 1.5 Hz, 1H), 7.46 – 7.32 (m, 1H), 7.04 (d, *J* = 10.2 Hz, 1H), 6.58 (d, *J* = 8.2 Hz, 1H), 5.77 (d, *J* = 10.2 Hz, 1H), 5.30 (s, 1H), 3.98 (s, 3H), 2.82 (s, 3H), 1.38 (s, 3H), 1.26 (s, 3H); Analytical HPLC: $t_R$ = 4.78 min, purity 97%, 5–95% $CH_3CN/H_2O$, gradient with constant 0.1% formic acid additive, 8 min run, 0.6 mL/min flow, UV detection at 254 nm; HRMS (ESI) calculated for $[M+H]^+$ 430.1649, found 430.1649.

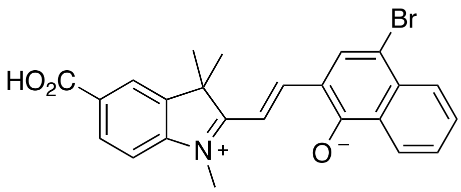

**6-bromo-1',3',3'-trimethylspiro[benzo[h]chromene-2,2'-indoline]-5'-carboxylic acid (4):** Compound **S4** (143 mg, 0.571 mmol, 1.0 eq) and compound **S6** (199 mg, 0.577 mmol, 1.0 eq) were suspended in acetonitrile (5 mL), and piperidine (0.113 mL, 1.14 mmol, 2.0 eq) was added. The mixture was heated at 82°C for 2 days. After cooling to room temperature, it was evaporated to dryness. The crude product was purified by column chromatography (0–50% EtOAc/cyclohexane) to afford the title compound as a dark purple solid (78 mg, 30%).$^1H$ NMR (400 MHz, $(CD_3)_2SO$) δ 8.00 (d, *J* = 8.5 Hz, 1H), 7.90 – 7.78 (m, 2H), 7.77 – 7.68 (m, 2H), 7.67 – 7.58 (m, 1H), 7.48 (ddd, *J* = 8.3, 6.8, 1.2 Hz, 1H), 7.19 (d, *J* = 10.2 Hz, 1H), 6.71 (d, *J* = 8.2 Hz, 1H), 5.91 (d, *J* = 10.2 Hz, 1H), 2.74 (s, 3H), 1.28 (s, 3H), 1.17 (s, 3H); Analytical HPLC: $t_R$ = 5.31 min, purity 95%, 5–95%

$CH_3CN/H_2O$, gradient with constant 0.1% formic acid additive, 8 min run, 0.6 mL/min flow, UV detection at 254 nm; HRMS (ESI) calculated for [M+H]+ 450.0699, found 450.0700.

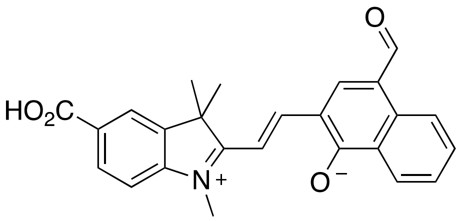

**6-formyl-1',3',3'-trimethylspiro[benzo[h]chromene-2,2'-indoline]-5'-carboxylic acid (5):** Compound **S5** (201 mg, 1.00 mmol, 1.0 eq), compound **S6** (346 mg, 1.00 mmol, 1.0 eq) were suspended in acetronitrile (10 mL), and piperidine (0.195 mL, 1.97 mmol, 2.0 eq) was added. The mixture was heated at 82°C for 2 days. After cooling to room temperature, it was evaporated to dryness. The crude product was purified by column chromatography (0–100% EtOAc/cyclohexane) to afford the title compound as a dark purple solid (143 mg, 36%). A small fraction was purified by HPLC for spectroscopic measurements. $^1H$ NMR (400 MHz, 1,4-dioxane-d8) δ 10.18 (s, 1H), 9.17 (d, *J* = 8.6 Hz, 1H), 7.92 (dd, *J* = 8.2, 1.7 Hz, 1H), 7.88 (d, *J* = 8.3 Hz, 1H), 7.85 (s, 1H), 7.78 (d, *J* = 1.7 Hz, 1H), 7.61 (ddd, *J* = 8.4, 6.8, 1.4 Hz, 1H), 7.42 (ddd, *J* = 8.3, 6.8, 1.2 Hz, 1H), 7.17 (d, *J* = 10.2 Hz, 1H), 6.65 (d, *J* = 8.2 Hz, 1H), 5.89 (d, *J* = 10.2 Hz, 1H), 2.79 (s, 3H), 1.36 (s, 3H), 1.23 (s, 3H); Analytical HPLC: $t_R$ = 3.70 min, purity 98%, 5–95% $CH_3CN/H_2O$, gradient with constant 0.1% formic acid additive, 8 min run, 0.6 mL/min flow, UV detection at 254 nm; HRMS (ESI) calculated for $[M+H]^+$ 400.1543, found 400.1544.

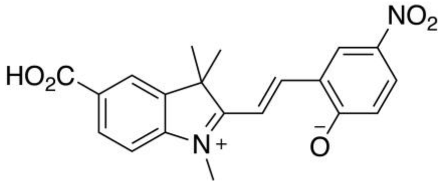

**(*E*)-2-(2-(5-carboxy-1,3,3-trimethyl-3*H*-indol-1-ium-2-yl)vinyl)-4-nitrophenolate (6):** synthesized according to published procedure.[8]

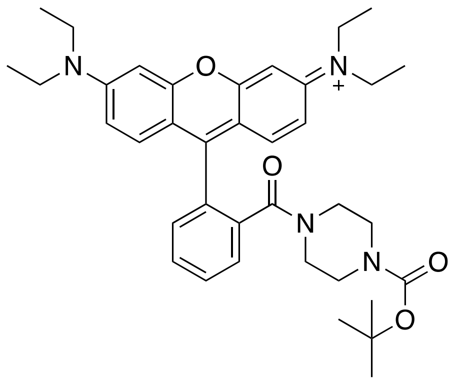

***N*-(9-(2-(4-(*tert*-butoxycarbonyl)piperazine-1-carbonyl)phenyl)-6-(diethylamino)-3*H*-xanthen-3-ylidene)-*N*-ethylethanaminium (Rh)**: Rhodamine B hydrochloride (500 mg, 1.04 mmol, 1 eq), 1-Boc piperazine (292 mg, 1.6 mmol, 1.5 eq) and HATU (790 mg, 2.1 mmol, 2.0 eq) were introduced in a vial. The vial was sealed under argon and DMF (30 mL) followed by DIEA (900 µL, 5.2 mmol, 5 eq) were added. The mixture was stirred at room temperature under argon for 18h. A solution of saturated $NH_4Cl$ was added, and the mixture was extracted with EtOAc (3x). The combined organic layers were washed with brine and $H_2O$, dried over anhydrous

$Na_2SO_4$, filtered and evaporated to dryness. The crude material was recrystallized in EtOH to afford the title compound as a pink solid (390 mg, 61%). $^1$H NMR (400 MHz, $CD_3OD$) δ 7.78 – 7.75 (m, 2H), 7.70 – 7.66 (m, 1H), 7.55 – 7.49 (m, 1H), 7.28 (d, *J* = 9.5 Hz, 2H), 7.08 (dd, *J* = 9.5, 2.4 Hz, 2H), 6.97 (d, *J* = 2.4 Hz, 2H), 3.69 (q, *J* = 7.2 Hz, 8H), 3.43 – 3.34 (m, 4H), 3.23 (s, 4H), 1.42 (s, 9H), 1.31 (t, *J* = 7.1 Hz, 12H); $^{13}$C NMR (100 MHz, $CD_3OD$) δ 169.6, 159.3, 157.2, 157.1, 156.0, 136.6, 133.2, 132.3, 131.7, 131.3, 131.3, 128.9, 115.4, 114.9, 97.3, 81.8, 46.9, 28.5, 12.8. (*Note:* $^{13}$C signals for the piperazine $CH_2$ groups are not visible on the NMR spectrum); HRMS (ESI) calculated for $[M+H]^+$ 611.3592, found 611.3588.

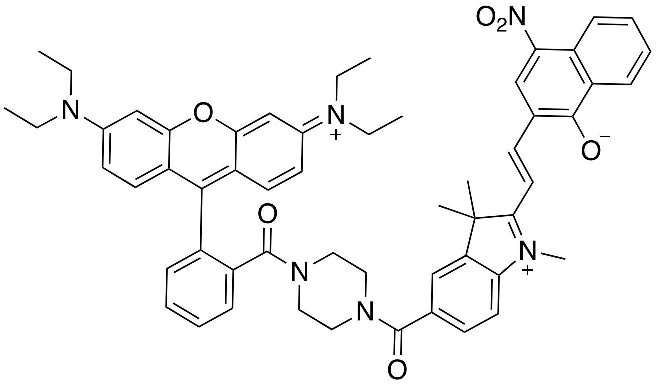

**(Rh-1)**: Compound **Rh** (300 mg, 0.49 mmol) was dissolved in $CH_2Cl_2$ (10 mL) and TFA (1 mL) was added. The mixture was stirred at room temperature for 1 hour. The mixture was evaporated to dryness and co-evaporated with toluene (3x), to afford the title compound as TFA salt in quantitative yield. The intermediate obtained (TFA salt, 45 mg, 72 µmol, 1.5 eq), compound **1** (20 mg, 48 µmol), and HATU (36 mg, 96 µmol, 2.0 eq) were introduced in a vial which was sealed under argon. DMF (2 mL) and DIEA (90 µL, 0.50 mmol, 10 eq) were added, and the mixture was stirred at room temperature under argon for 30 minutes. The mixture was evaporated to dryness and purified by column chromatography (0–10% MeOH/$CH_2Cl_2$) followed by preparative HPLC (10–95% MeCN/$H_2O$) to afford the title compound as a dark purple solid (10 mg, 19%).

$^1$H NMR (400 MHz, $(CD_3)_2SO$ containing 5 mM DMF. *Note:* in these conditions the MC form is present at >85% and only the signals corresponding to this isomer are reported. Integration on the spectrum is however in some instances higher due to the contribution of the SP isomer) δ 8.94 (s, 1H), 8.77 (dd, *J* = 4.4, 1.4 Hz, 1H), 8.60 (d, *J* = 8.6 Hz, 1H), 8.54 (dd, *J* = 8.4, 1.4 Hz, 1H), 8.35 (d, *J* = 7.9 Hz, 1H), 8.27 (br s, 1H), 7.79 – 7.68 (m, 4H), 7.63 (d, *J* = 8.3 Hz, 1H), 7.56 – 7.44 (m, 3H), 7.22 – 7.05 (m, 4H), 6.96 (d, *J* = 2.3 Hz, 3H), 3.71 – 3.58 (m, 8H), 3.52 – 3.47 (m, 4H), 3.20 – 3.08 (m, 4H), 2.73 (s, 3H), 1.75 (s, 6H), 1.27 – 1.19 (m, 12H); Analytical HPLC: $t_R$ = 3.95 min, purity 98%, 5–95% $CH_3CN/H_2O$, gradient with constant 0.1% formic acid additive, 8 min run, 0.6 mL/min flow, UV detection at 254 nm; HRMS (ESI) calculated for $[M+H]^+$ 909.4334, found 909.4329.

### NMR Spectra and HPLC Traces

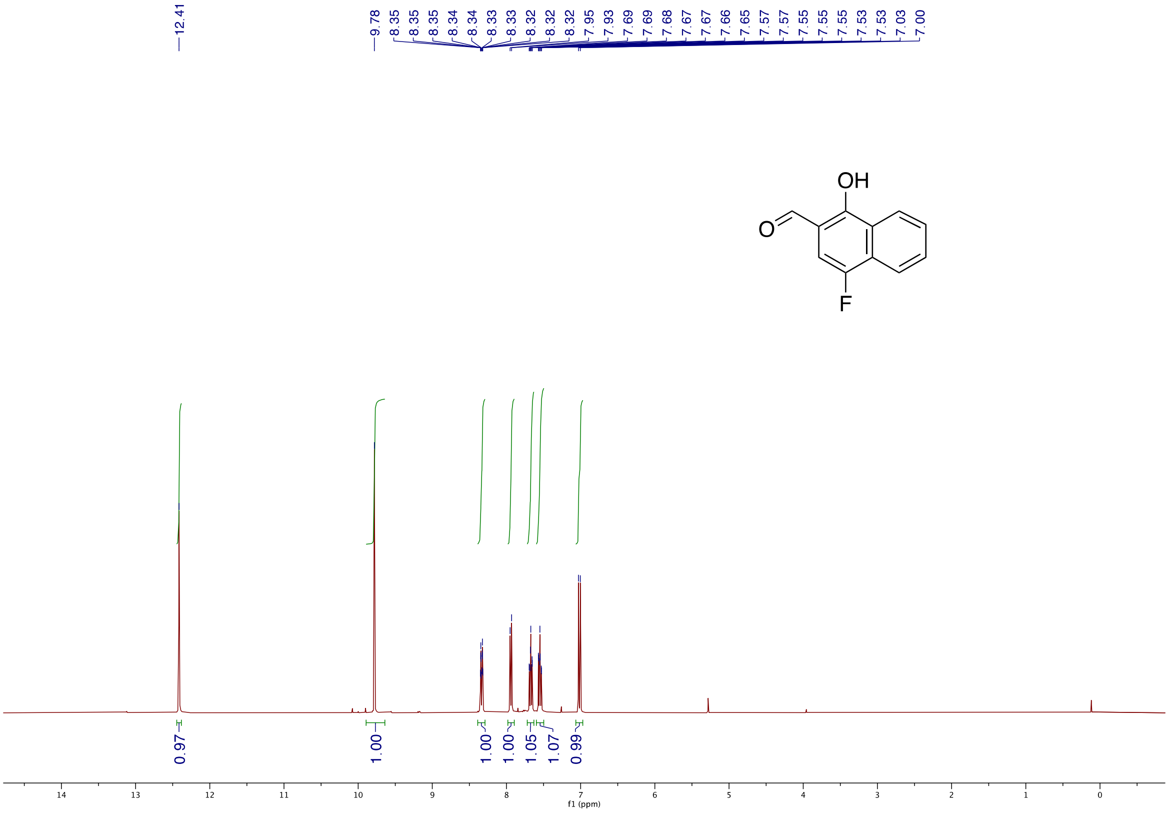

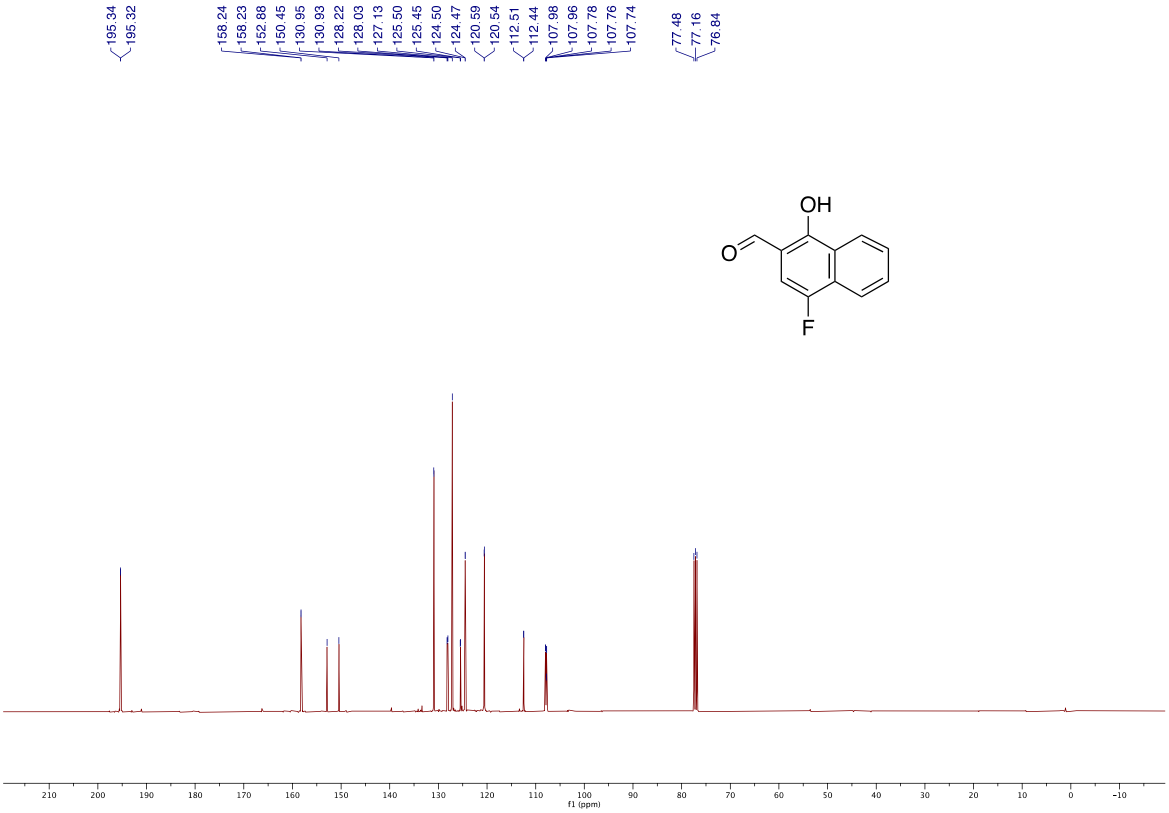

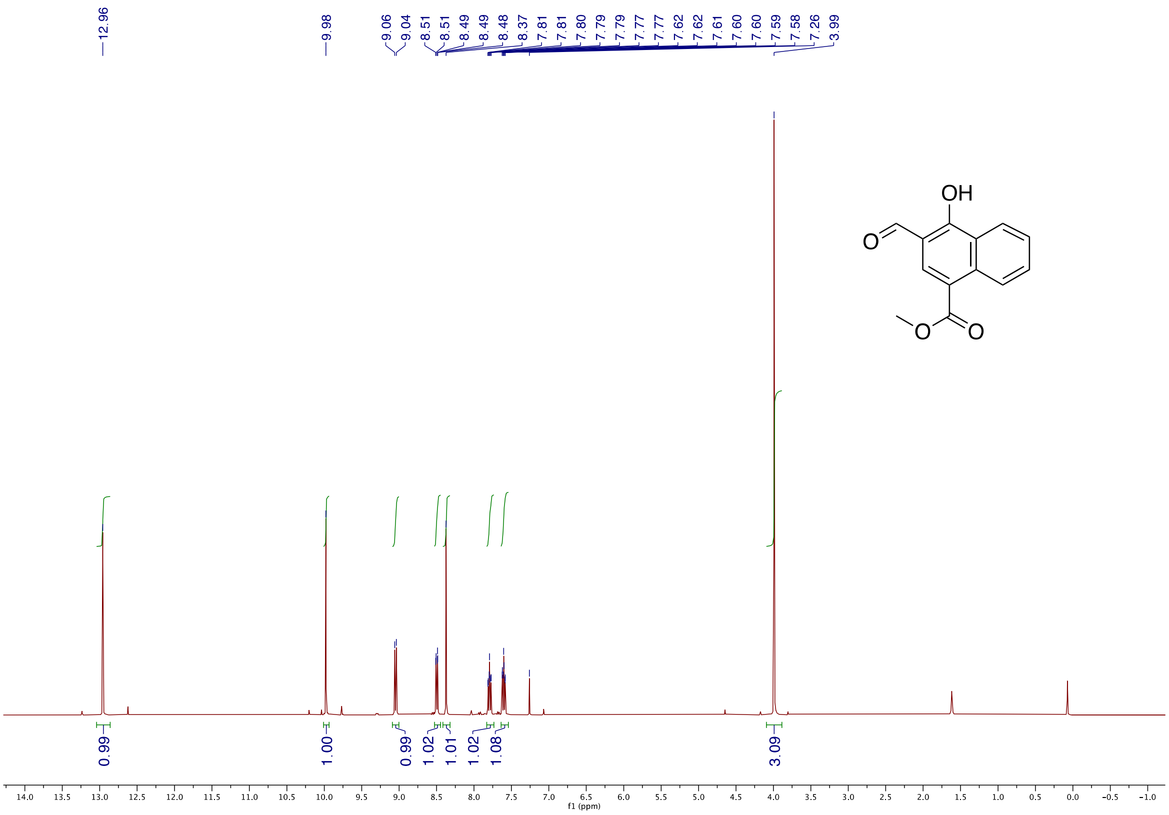
12.96
9.98
9.06
9.04
8.51
8.51
8.49
8.49
8.48
8.37
7.81
7.81
7.80
7.79
7.79
7.77
7.77
7.62
7.62
7.61
7.60
7.60
7.59
7.58
7.26
3.99
OH
O
O
O
0.99
1.00
0.99
1.02
1.01
1.02
1.08
3.09
f1 (ppm)

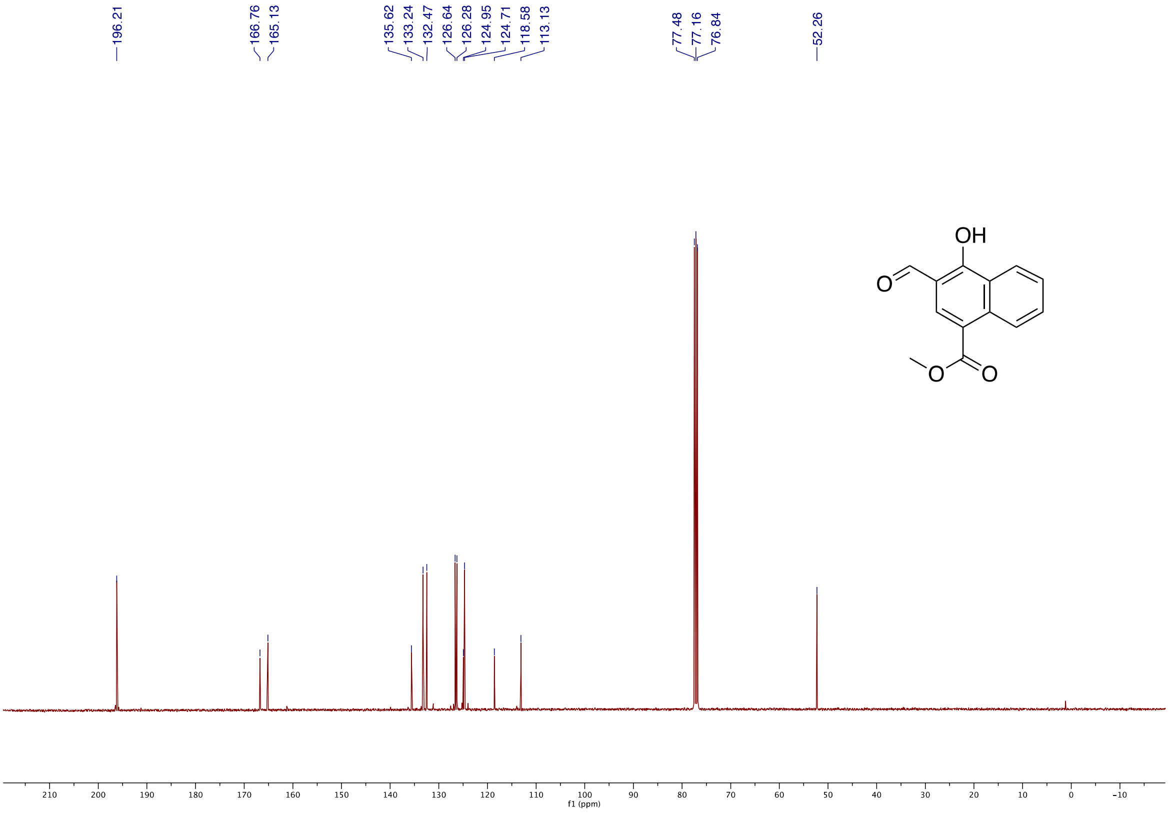
196.21
166.76
165.13
135.62
133.24
132.47
126.64
126.28
124.95
124.71
118.58
113.13
77.48
77.16
76.84
52.26
OH
O
O
O
f1 (ppm)

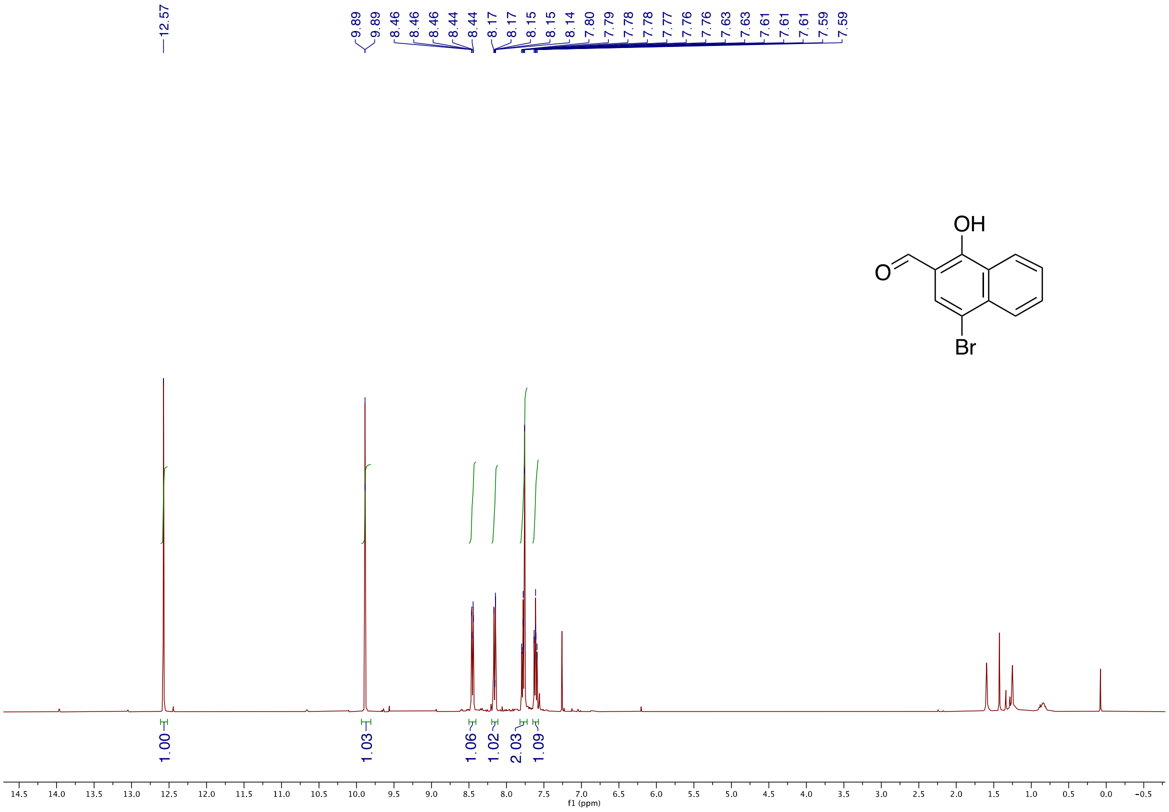
12.57
9.89
9.89
8.46
8.46
8.46
8.44
8.44
8.17
8.17
8.15
8.15
8.14
7.80
7.79
7.78
7.78
7.77
7.76
7.76
7.63
7.63
7.61
7.61
7.61
7.59
7.59
OH
O
Br
1.00
1.03
1.06
1.02
2.03
1.09
14.5
14.0
13.5
13.0
12.5
12.0
11.5
11.0
10.5
10.0
9.5
9.0
8.5
8.0
7.5
7.0
6.5
6.0
5.5
5.0
4.5
4.0
3.5
3.0
2.5
2.0
1.5
1.0
0.5
0.0
-0.5
f1 (ppm)

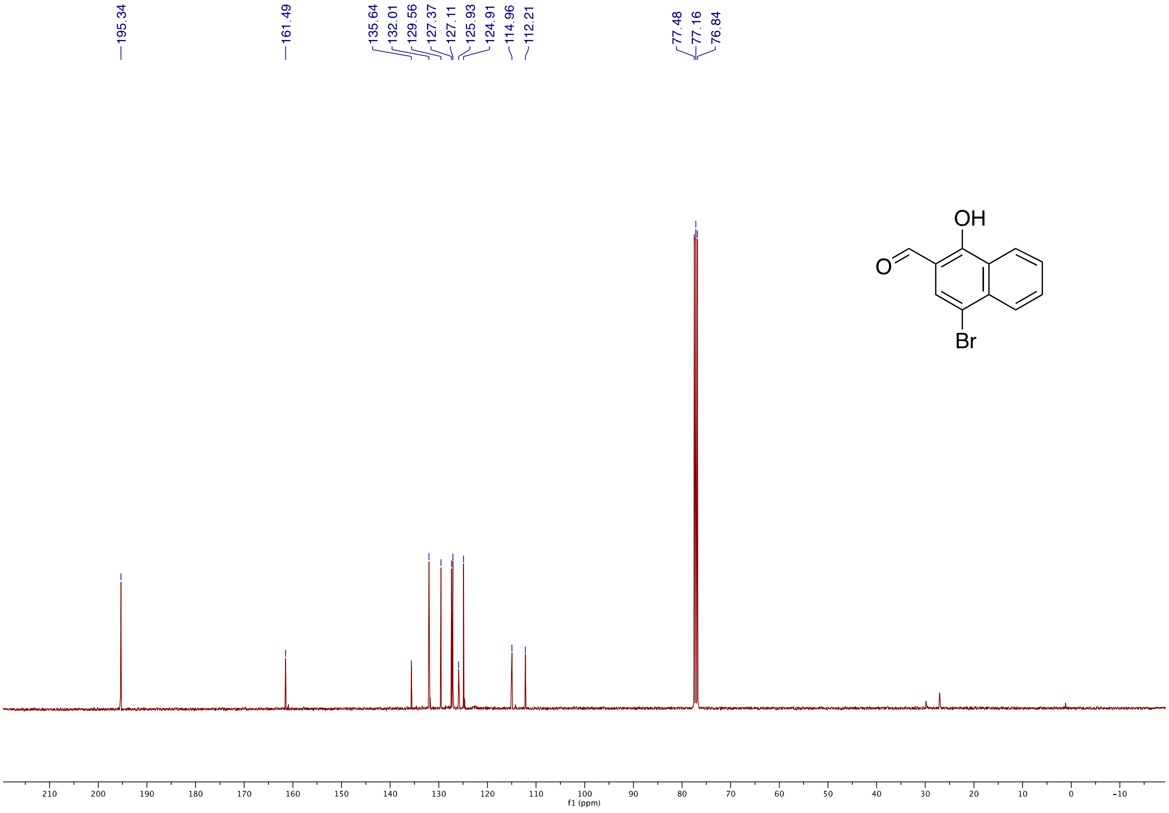
195.34
161.49
135.64
132.01
129.56
127.37
127.11
125.93
124.91
114.96
112.21
77.48
77.16
76.84
OH
O
Br
210
200
190
180
170
160
150
140
130
120
110
100
90
80
70
60
50
40
30
20
10
0
-10
f1 (ppm)

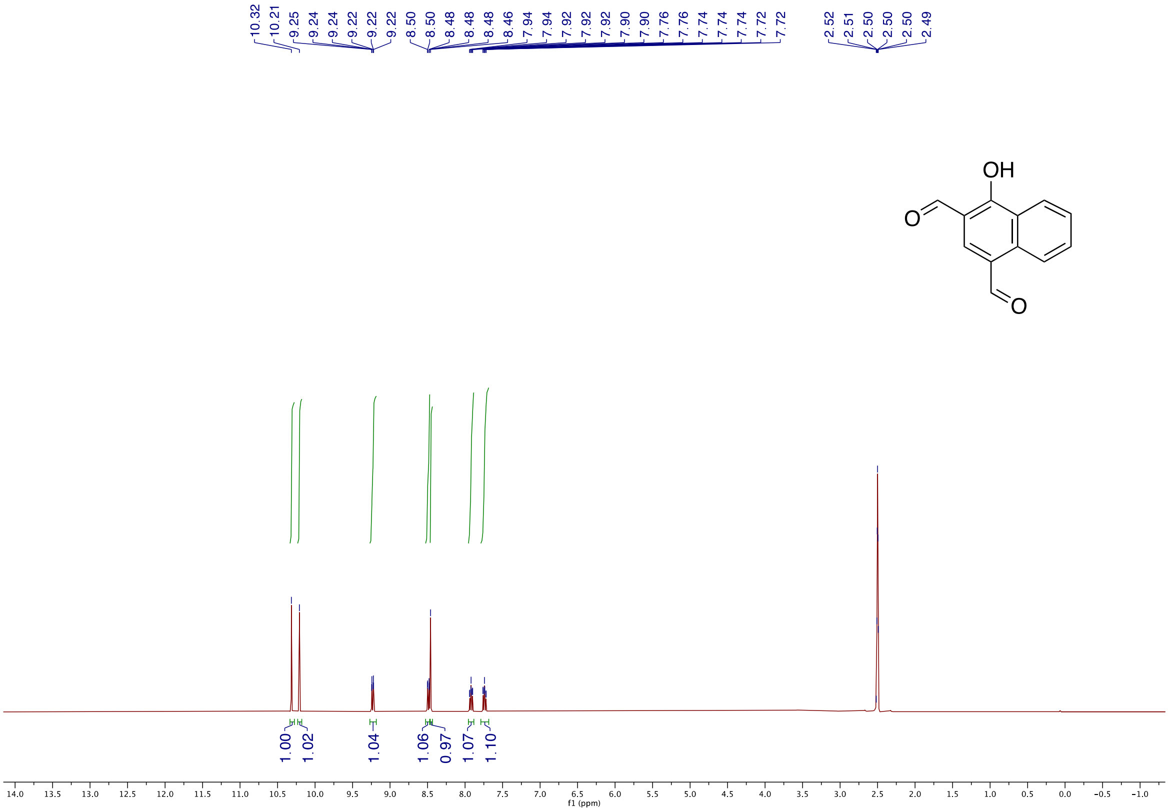
10.32
10.21
9.25
9.24
9.24
9.22
9.22
9.22
8.50
8.50
8.48
8.48
8.48
8.46
7.94
7.94
7.92
7.92
7.92
7.90
7.90
7.76
7.74
7.74
7.74
7.72
7.72
2.52
2.51
2.50
2.50
2.50
2.49
OH
O
O
1.00
1.02
1.04
1.06
0.97
1.07
1.10
f1 (ppm)

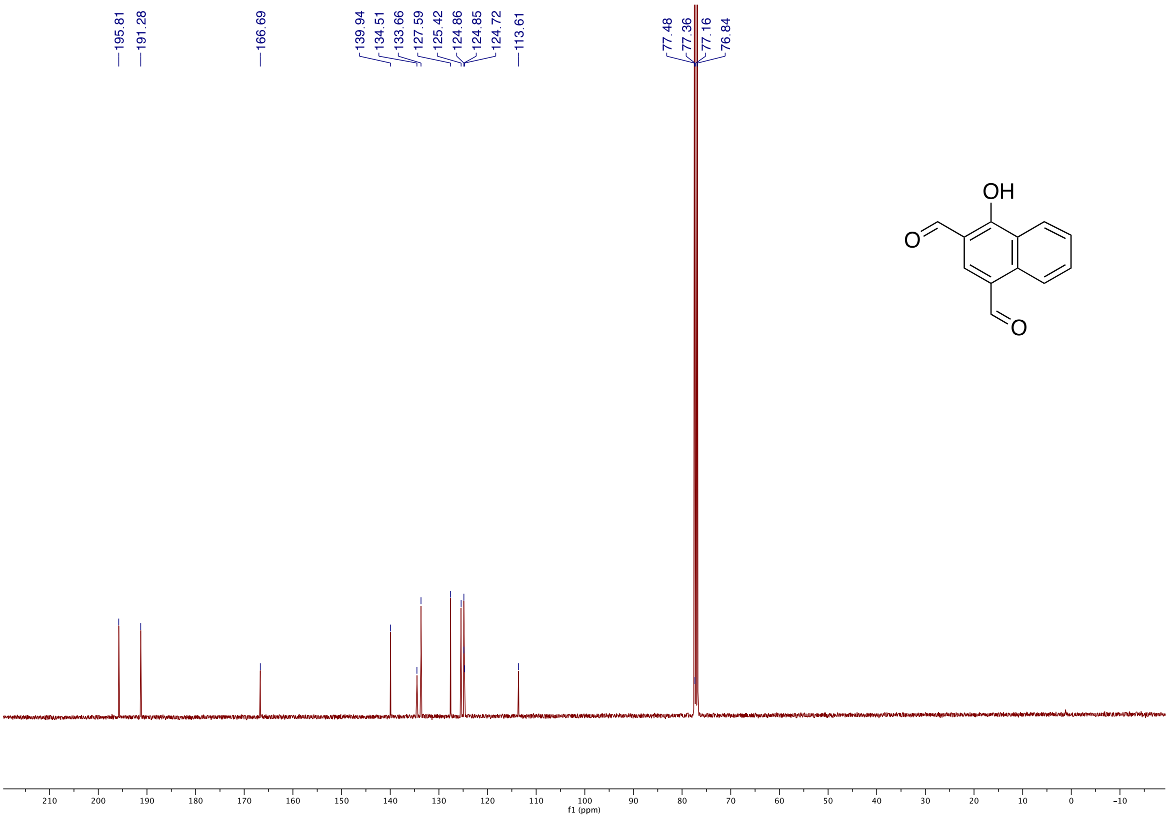
195.81
191.28
166.69
139.94
134.51
133.66
127.59
125.42
124.86
124.85
124.72
113.61
77.48
77.36
77.16
76.84
OH
O
O
f1 (ppm)

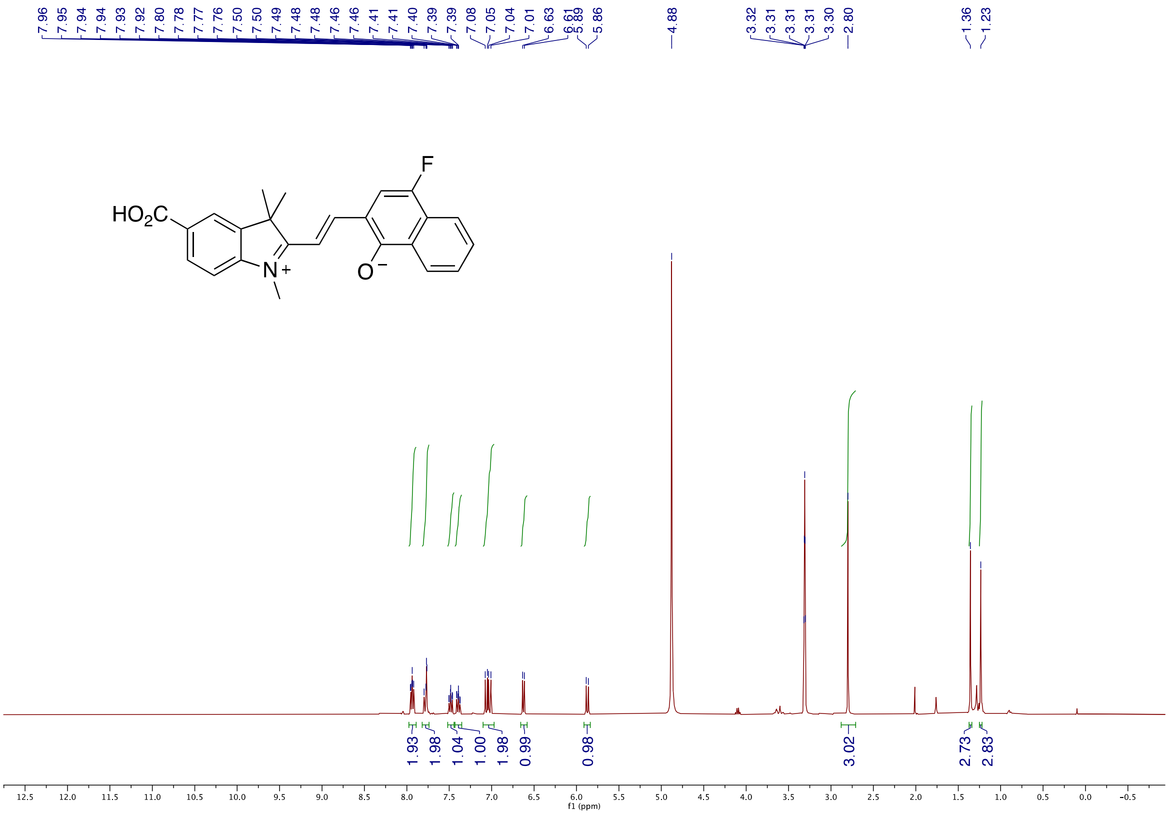
7.96 7.95 7.94 7.94 7.93 7.92 7.80 7.78 7.77 7.76 7.50 7.50 7.49 7.48 7.48 7.46 7.41 7.41 7.40 7.39 7.39 7.08 7.05 7.04 7.01 6.63 6.61 5.89 5.86
4.88
3.32 3.31 3.31 3.31 3.30 2.80
1.36 1.23
HO2C
N+
O−
F
1.93 1.98 1.04 1.00 1.98 0.99 0.98
3.02
2.73 2.83
f1 (ppm)

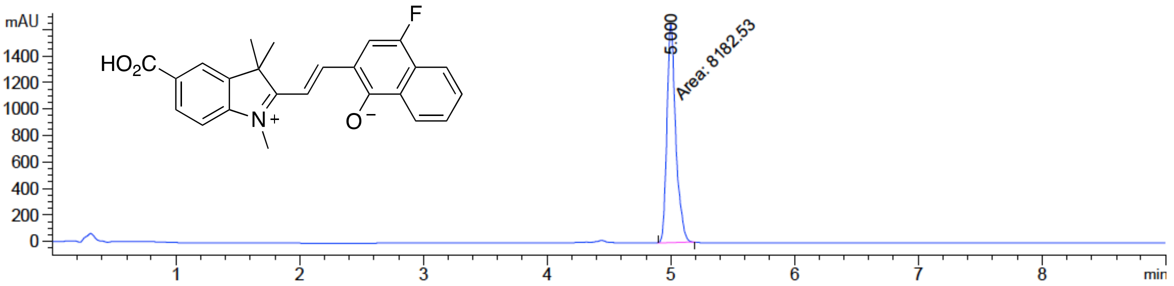
mAU
1400
1200
1000
800
600
400
200
0
5.000
Area: 8182.53
HO2C
N+
O−
F
1 2 3 4 5 6 7 8
min

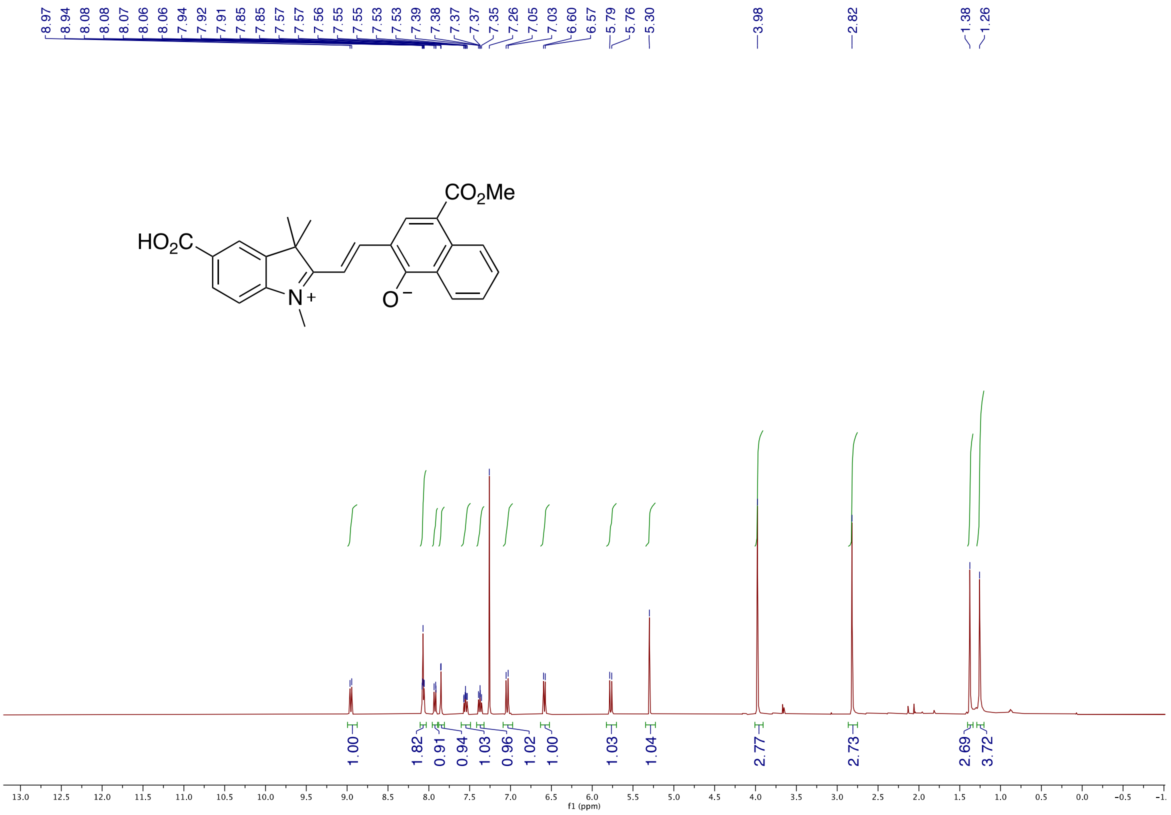
8.97
8.94
8.08
8.08
8.07
8.06
8.06
7.94
7.92
7.91
7.85
7.85
7.57
7.57
7.56
7.55
7.55
7.53
7.53
7.39
7.38
7.37
7.37
7.35
7.26
7.05
7.03
6.60
6.57
5.79
5.76
5.30
3.98
2.82
1.38
1.26
$CO_2Me$
$HO_2C$
$N^+$
$O^-$
1.00
1.82
0.91
0.94
1.03
0.96
1.02
1.00
1.03
1.04
2.77
2.73
2.69
3.72
f1 (ppm)

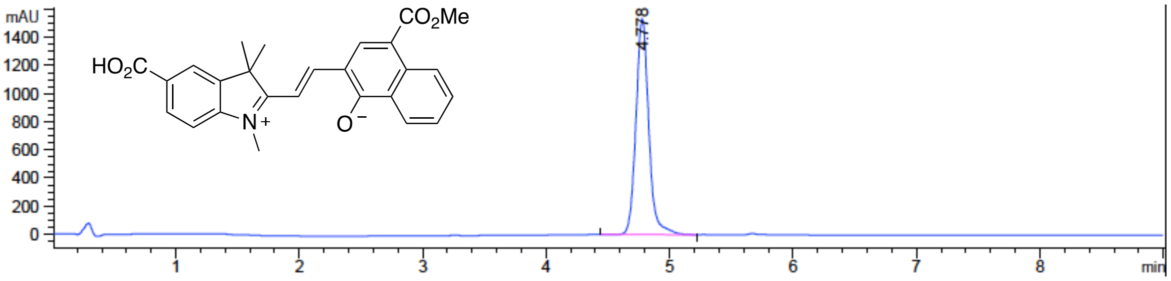
mAU
1400
1200
1000
800
600
400
200
0
$CO_2Me$
$HO_2C$
$N^+$
$O^-$
4.778
1
2
3
4
5
6
7
8
min

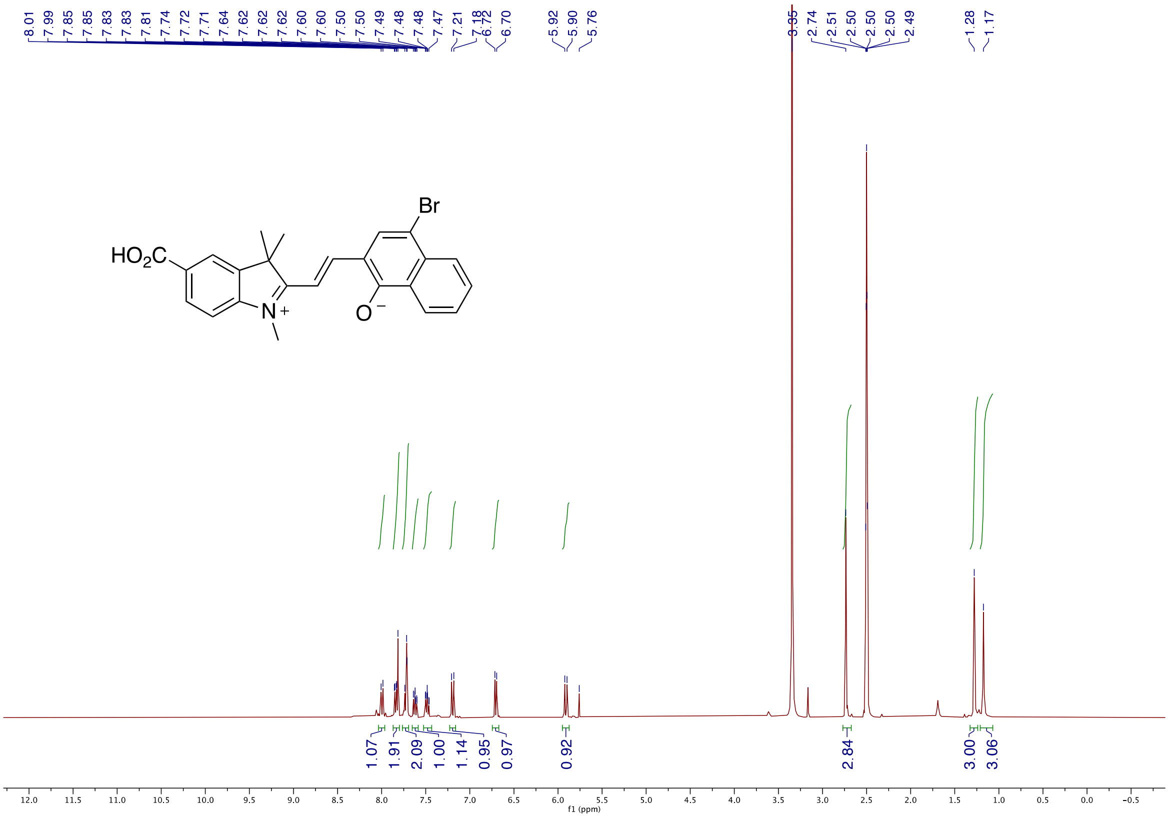

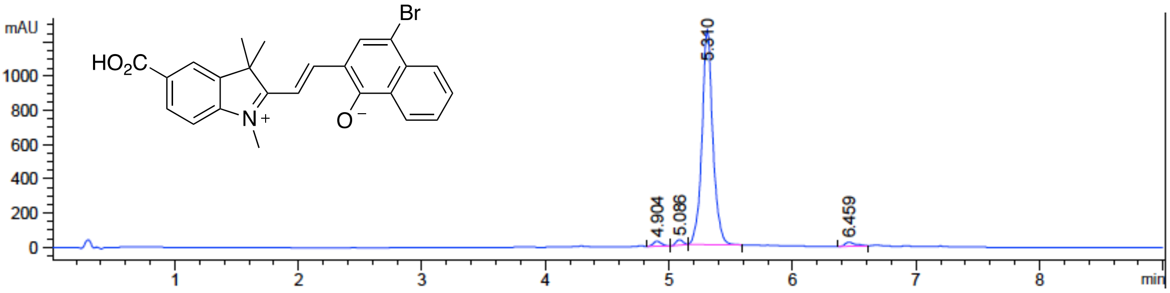

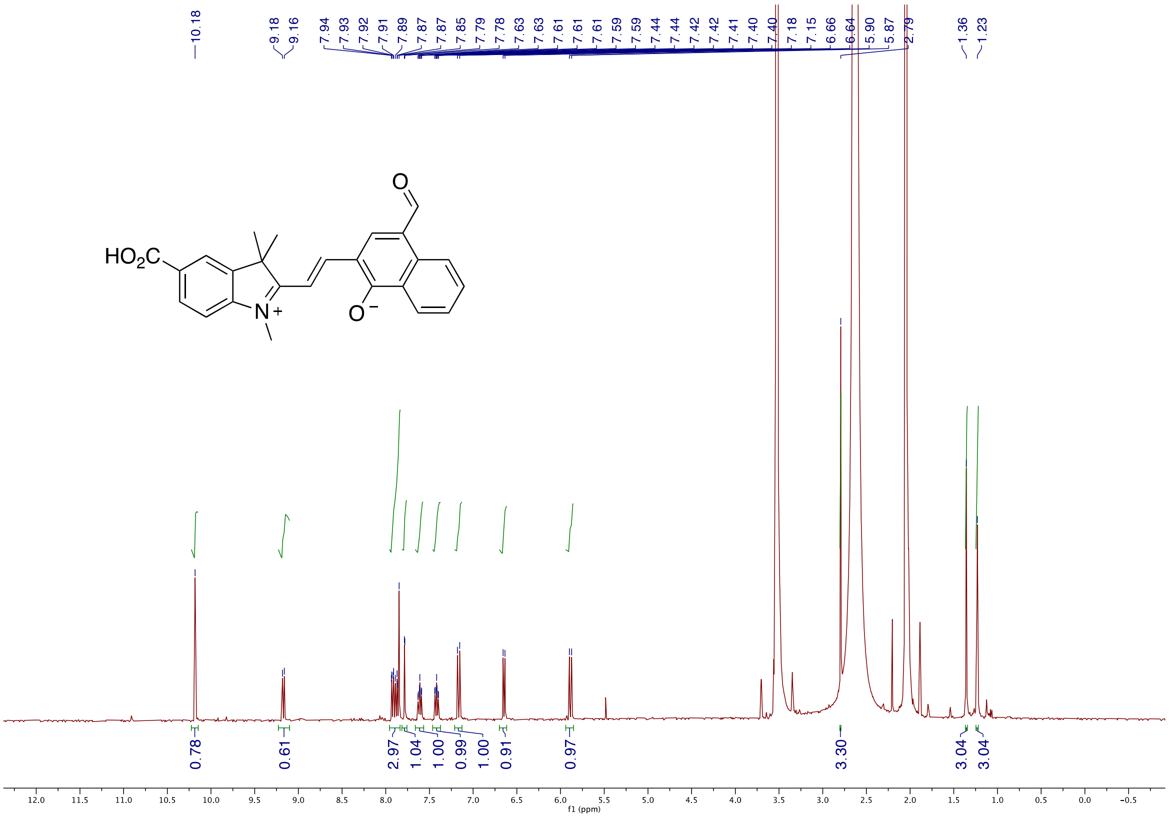

HO2C
N+
O
O−
f1 (ppm)

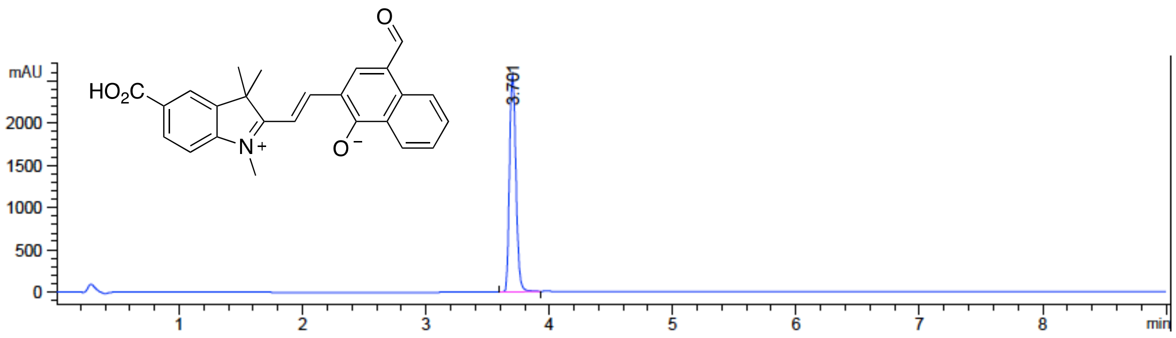

mAU
HO2C
N+
O
O−
3.701
min

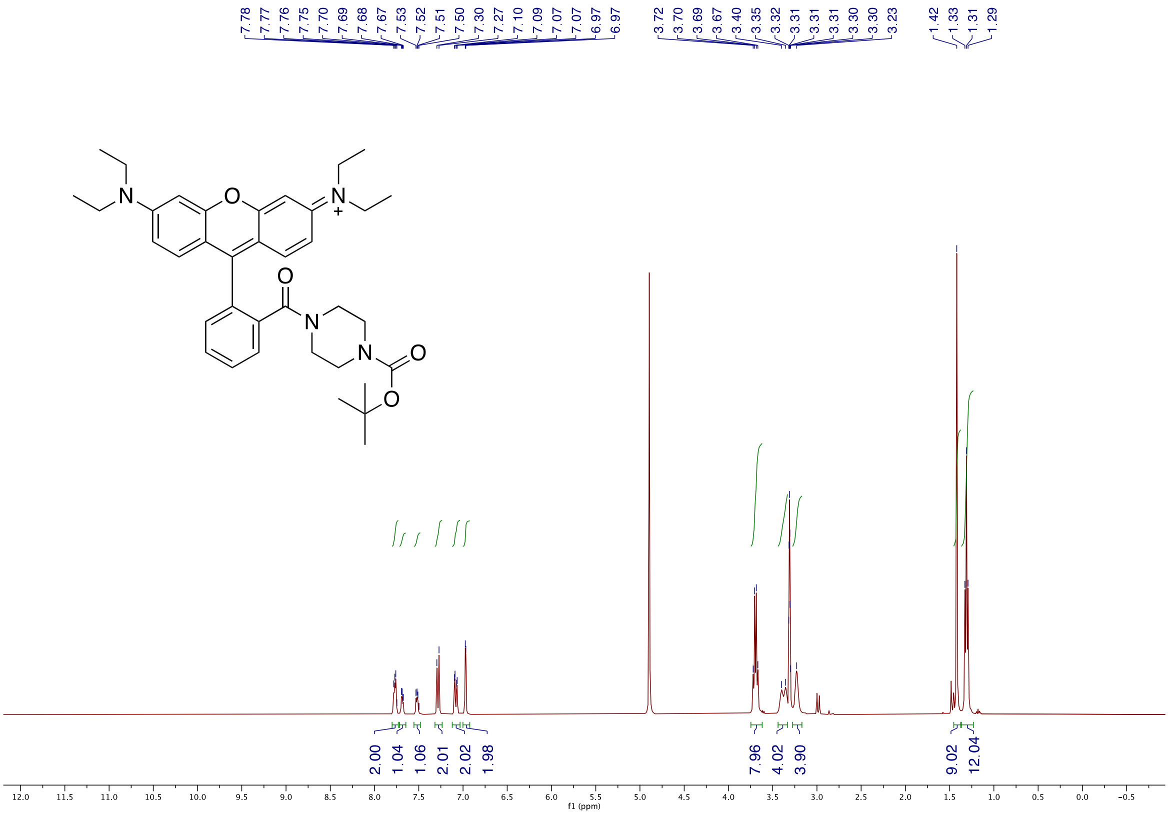
7.78 7.77 7.76 7.75 7.70 7.69 7.68 7.67 7.53 7.52 7.51 7.50 7.30 7.27 7.10 7.09 7.07 7.07 6.97 6.97
3.72 3.70 3.69 3.67 3.40 3.35 3.32 3.31 3.31 3.31 3.30 3.30 3.23
1.42 1.33 1.31 1.29
2.00 1.04 1.06 2.01 2.02 1.98
7.96 4.02 3.90
9.02 12.04
f1 (ppm)

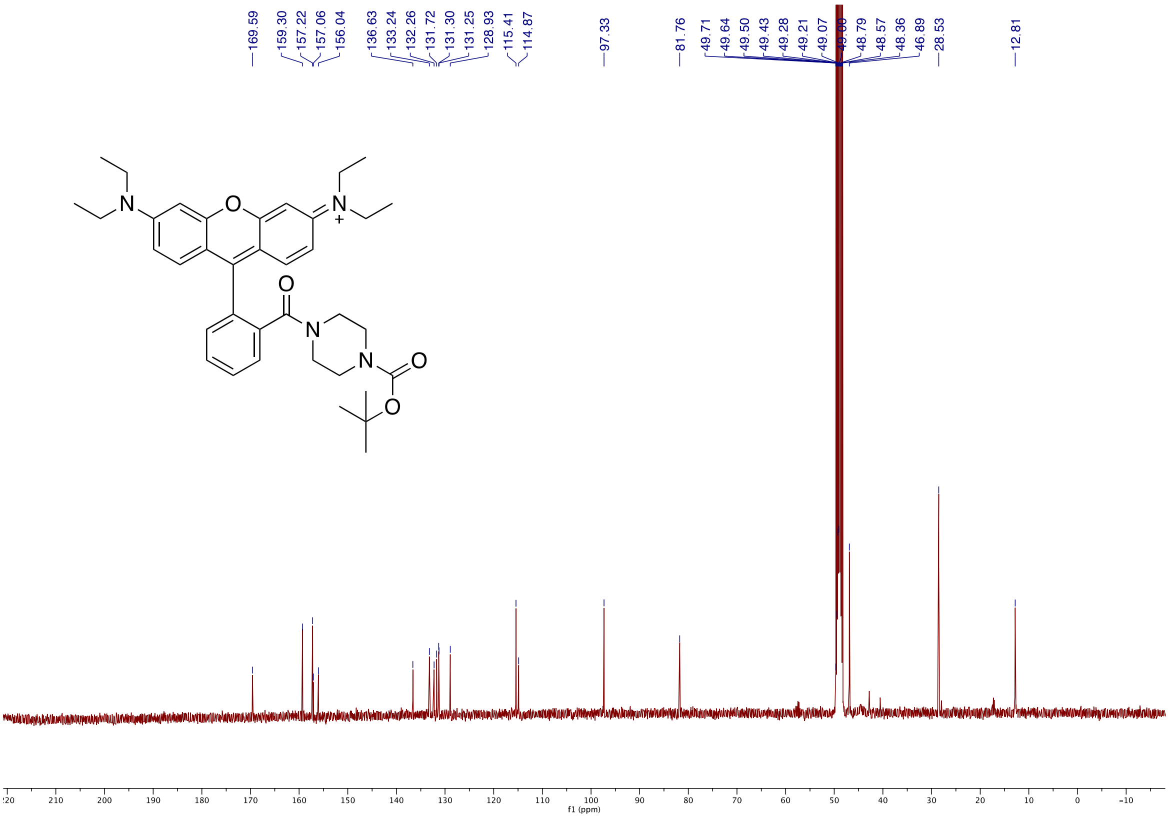
169.59
159.30 157.22 157.06 156.04
136.63 133.24 132.26 131.72 131.30 131.25 128.93
115.41 114.87
97.33
81.76
49.71 49.64 49.50 49.43 49.28 49.21 49.07 48.79 48.57 48.36 46.89
28.53
12.81
f1 (ppm)

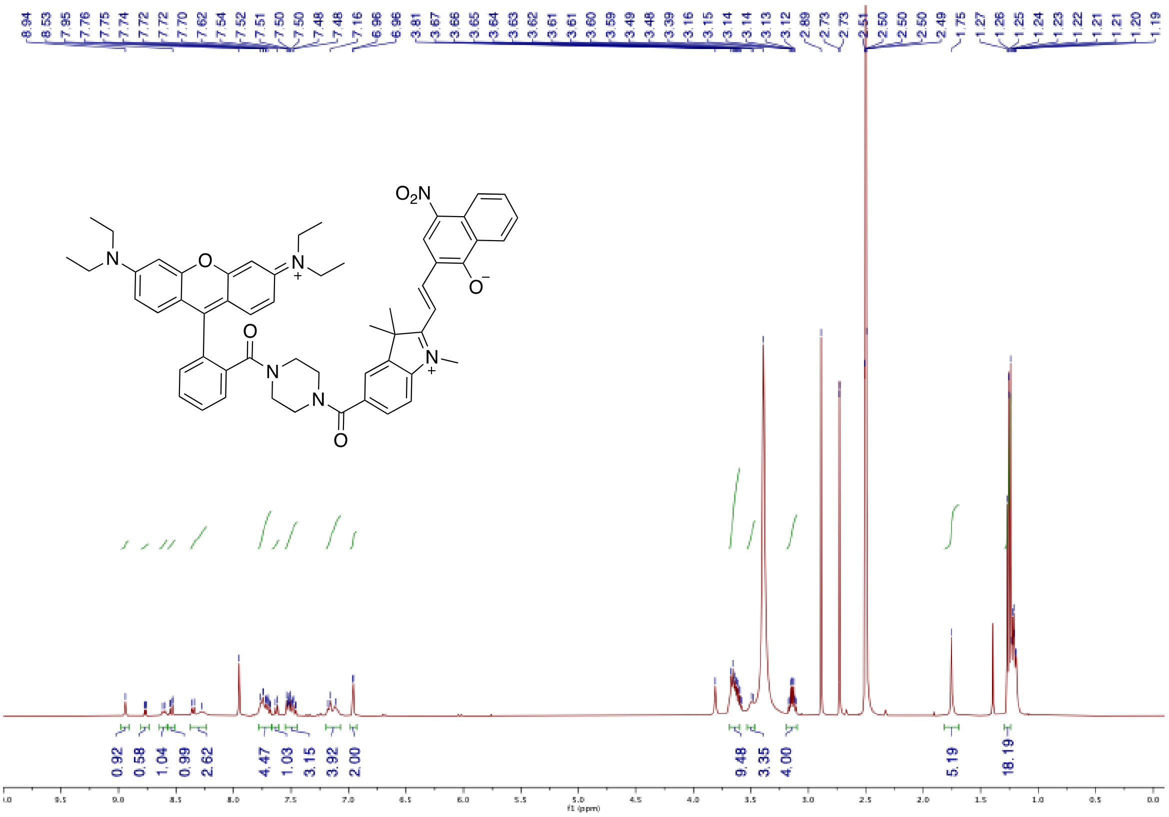
8.94 8.53 7.95 7.76 7.75 7.74 7.72 7.72 7.70 7.62 7.54 7.52 7.51 7.50 7.50 7.48 7.48 7.16 6.96 6.96 3.81 3.67 3.66 3.65 3.64 3.63 3.62 3.61 3.61 3.60 3.59 3.49 3.48 3.39 3.16 3.15 3.14 3.14 3.13 3.12 2.89 2.73 2.73 2.51 2.50 2.50 2.50 2.49 1.75 1.27 1.26 1.25 1.24 1.23 1.22 1.21 1.21 1.20 1.19
$O_2N$
N
O
N
+
O
N
N
O
N
+
O
0.92 0.58 1.04 0.99 2.62 4.47 1.03 3.15 3.92 2.00 9.48 3.35 4.00 5.19 18.19
f1 (ppm)

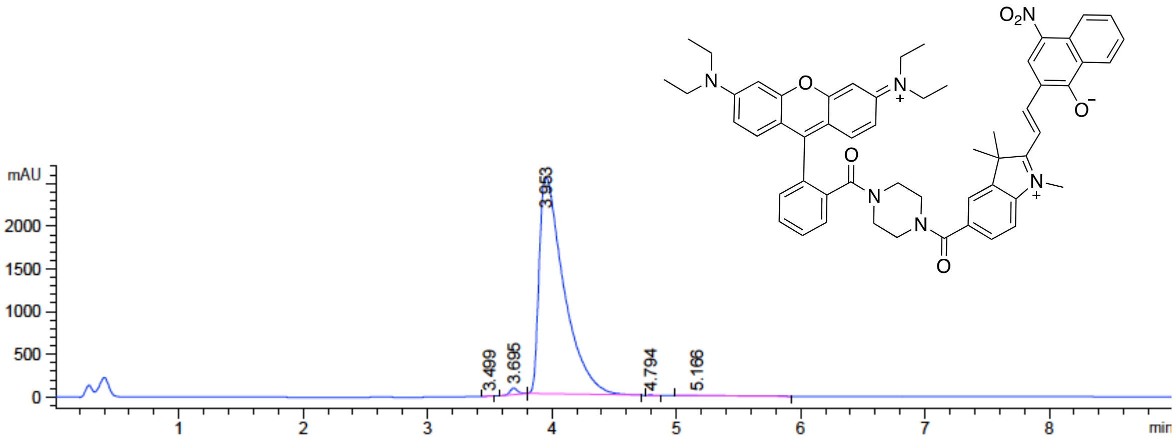
mAU
2000
1500
1000
500
0
3.499
3.695
3.953
4.794
5.166
1 2 3 4 5 6 7 8
min
$O_2N$
N
O
N
+
O
N
N
O
N
+
O
